# Earth-Observation and Environmental Vision Transformers Reveal Genome–Environment Associations in Macroalgae

**DOI:** 10.64898/2025.12.30.696986

**Authors:** Alexandra Mystikou, David R. Nelson, Diana C. El Assal, Ashish K. Jaiswal, Mehar Sultana, Cecilia Rad-Menendez, Noura Al-Mansoori, Sewar T. Elias, Ma-sum Abdul-Hamid, Layanne Nayfeh, Nizar Drou, John A. Burt, David H. Green, Kourosh Salehi-Ashtiani

## Abstract

Macroalgae thrive in extreme environments, yet the genomic basis of their tolerance remains poorly resolved. We describe nine Arabian Gulf macroalgae and integrate them with 117 published genomes (126 total; 70 Rhodophyta, 43 Ochrophyta, 13 Chlorophyta) to test genome–environment associations. Google Earth Engine (GEE) for broad-scale oceanography and 10-meter resolution AlphaEarth Foundations (AEF) embeddings for fine-scale habitat heterogeneity. We identified 157 significant (FDR q < 0.05) correlations with global GEE variables—including a strong negative temperature association with DUF3570—while AEF embeddings uncovered over 1,000 lineage-specific signals within Rhodophyta and identified climate-driven Pfam modules. The von Willebrand factor type-A domain emerged as uniquely robust across all frameworks and enriched in Arabian Gulf species. In the Arabian Gulf, enrichment of this domain is consistent with selection for adhesion mechanisms capable of withstanding chronic hydrodynamic stress compounded by high temperature and salinity. These results demonstrate that converging remote sensing with deep learning identifies conserved and lineage-specific genomic signatures of ecological differentiation across diverse macroalgal lineages.

## INTRODUCTION

Marine algae are foundational organisms in ocean ecosystems, contributing approximately 50% of Earth’s oxygen production through photosynthesis and serving as the base of marine food webs.^1^ These multicellular photosynthetic organisms—classified as brown (Ochrophyta), green (Chlorophyta), and red (Rhodophyta) algae based on their photosynthetic pigments—have colonized virtually every aquatic habitat on Earth. From the ice-covered waters of polar regions to tropical coral reefs, from intertidal zones exposed to open air for hours to depths exceeding 268 meters where less than 0.1% of surface light penetrates,^2^ representatives from the three macroalgal phyla embody environmental plasticity.

As climate change accelerates and human populations grow, macroalgae are increasingly recognized as sustainable resources for food security, pharmaceutical development, biomaterial production, and climate mitigation through CO2 sequestration.^3^ Large-scale cultivation projects, exemplified by kelp forest restoration initiatives,^4^ highlight their potential in carbon capture and habitat restoration. The Mediterranean invasion of *Caulerpa taxifolia*,^5,6^ the Pacific-to-Atlantic spread of *Gracilaria vermiculophylla*,^7^ and the transcontinental establishment of *Undaria pinnatifida*^8^ show how genomes that confer success in native habitats can drive invasive behavior^9^ in new environments. Despite their ecological importance and economic potential, our understanding of macroalgal genomics lags far behind that of terrestrial plants and other model organisms. Macroalgal genomics studies are limited to only a handful,^10–20^ but over 10,000 macroalgal species have been described. The current genomic shortfall continues to limit understanding of the molecular basis of environmental adaptation,^21^ constraining the ability to predict species’ responses to climate change or to leverage their traits for biotechnological applications.

As one of the world’s most thermally and osmotically extreme marine environments, the Arabian Gulf experiences summer temperatures exceeding 35°C^22^ and salinities exceeding 44 PSU^23–28^ — conditions that push marine life to physiological limits and may represent future ocean states in other regions under climate change.^29–31^ The macroalgae thriving in these extreme conditions for millennia potentially harbor genomic innovations for multi-stressor tolerance with applications in aquaculture, climate adaptation prediction, and understanding invasiveness. Identifying the genetic basis of environment associations in macroalgae would enable several practical advances: predicting species range shifts under ocean warming, selecting thermotolerant strains for aquaculture operations increasingly affected by marine heatwaves, and understanding why certain species (*Caulerpa, Sargassum, Undaria*) successfully invade novel environments while others do not.

We integrated nine newly described Arabian Gulf species with 117 global genomes (Table S1) to compile a master dataset to address these questions. Although draft assemblies of these nine UAE macroalgal genomes were previously deposited and included as public data in a large-scale comparative genomics study,^32^ their isolation, environmental context, and genomic characterization have not been formally described. Here, we provide the first dedicated description and analyses of these genomes, contextualized within all available macroalgae genomes.

We analyzed genome–environment relationships using two satellite-derived environmental characterizations: correlations with physically interpretable oceanographic variables extracted via Google Earth Engine (GEE) and learned environmental embeddings generated by vision transformers trained on petabyte-scale imagery (AEF). Together, these representations enable detection of genome–environment associations that extend beyond traditional single-variable analyses. By integrating interpretable oceanographic measurements with learned environmental structure, we test whether protein domain profiles predict environmental tolerance and identify candidate gene families associated with sustainable macroalgal colonization under a variety of environmental conditions.

## RESULTS

### Global macroalgal genome dataset spanning extreme environmental gradients

To investigate macroalgal genome-environment associations on a global scale, we started by expanding the sample size of lineages inhabiting less-well represented hypersaline subtropical habitats. We integrated the genome of nine species that we have isolated from the Arabian Gulf (Figure 1) with 117 publicly available macroalgal genomes spanning global environmental gradients (Table S1). The final dataset included 126 genomes representing all three major macroalgal phyla: 70 Rhodophyta (red algae), 43 Ochrophyta (brown algae), and 13 Chlorophyta (green algae), distributed across 80 genera, five climates from polar (n = 15) to tropical (n = 48), three salinity regimes (marine, brackish, freshwater), and depth ranges from intertidal to 268m subtidal.

**Figure 1.**
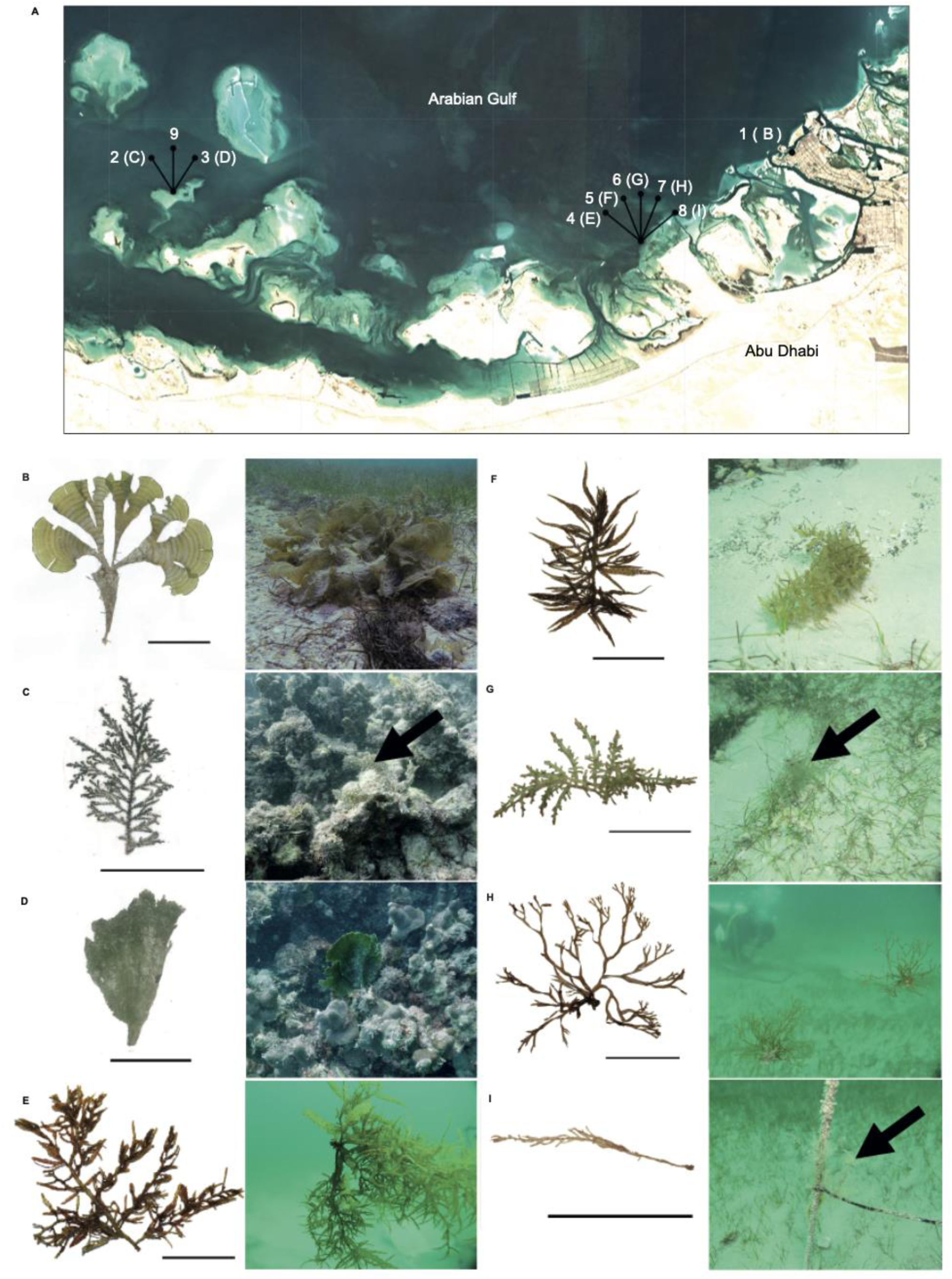
Geographic distribution and morphological diversity of sampled Arabian Gulf macroalgae. (A) Map showing collection sites of nine macroalgal species along the UAE coast. Sampling locations include: (1) North Corniche, Abu Dhabi (intertidal); (2-3, (9)) Al Hiel (subtidal, 3-7m depth, specimen 9 represents an endophyte of (D)); (4-7) Dhabiya (subtidal, 3-7m depth); (8) Ras Ghurab (subtidal, 5m depth). Collection sites were North Corniche, Al Hiel, Dhabiya, and Ras Ghurab. Numbers in parentheses correspond to specimen codes in panels B-I. (B-I) Morphological diversity of collected specimens. Left panels show herbarium specimens; right panels show in situ underwater photographs demonstrating natural habitat and growth forms. Scale bars, 5 cm. Species shown: (B) *Padina boergesenii* (NYAE_20191204_1), Ochrophyta. The rolled-blade morphology is typical of Dictyotales. (C) *Polycladia myrica* (NYAE_20191205_1), Ochrophyta. Black arrow indicates pneumatocysts (gas-filled flotation structures). (D) *Avrainvillea amadelpha* (NYAE_20191205_6a), Chlorophyta. This specimen hosted an endophytic brown alga (NYAE_20191205_6b). (E) *Sargassum latifolium* (NYAE_20200302_1), Ochrophyta. Representative of the dominant macroalgal genus in the Arabian Gulf. (F) *Sargassum angustifolium* (NYAE_20200302_2), Ochrophyta. Black arrow indicates characteristic narrow blade morphology distinguishing it from *S. latifolium*. (G) *Chondria dasyphylla* (NYAE_20200302_4), Rhodophyta. The only red algal representative collected, showing typical branching pattern. (H) *Canistrocarpus cervicornis* (NYAE_20200302_5), Ochrophyta. Specimen was epizoic on gastropod shell, indicating opportunistic substrate utilization. (I) Unidentified brown alga (NYAE_20200302_6), Ochrophyta. Black arrow indicates attachment to artificial substrate (monitoring station pole), suggesting recent colonization. See also Table S1 for detailed collection data and GPS coordinates. Figure 1: UAE macroalgal genomes sample locations, habitats and herbarium specimens.

The nine Arabian Gulf species provide the only available genomes from hypersaline, hyperthermal marine conditions predicted to become widespread under climate change.^33^ Field-collected UAE strains included six brown algae (*Padina boergesenii, Polycladia myrica, Sargassum latifolium, S. angustifolium, Canistrocarpus cervicornis*, and one unidentified endophyte), one red alga (*Chondria dasyphylla*), and one green alga (*Avrainvillea amadelpha*), sampled from four locations in Abu Dhabi(Al Hiel, Dhabiya, Ras Ghurab, Abu Dhabi) spanning intertidal and subtidal zones to 7m depth (Fig. 1A-I). Several species colonized recently bleached dead coral substrates, including *A. amadelpha*—documented as invasive in Mediterranean and Hawaiian waters.^34^

To identify protein domain families associated with environmental gradients, we performed exclusion analyses (Fig. 2) and correlated Pfam abundances with manually curated environmental metadata from published records (Fig. 3; Table S1). Spearman rank correlation analysis identified 377 significant protein domain–environment associations at p < 0.001 (n = 126 samples). After false discovery rate correction (Benjamini-Hochberg FDR), 114 correlations remained statistically robust (q < 0.05). Bonferroni correction for family-wise error rate (α = 0.05/41,852 tests = 1.2×10⁻⁶) yielded 10 protein domain families surviving genome-wide significance thresholds. Geographic coordinates had the most correlations (longitude: 150 associations, latitude: 55), indicating that protein domain distributions track biogeographic structure. Temperature associations were predominantly negative (133 correlations; 80 positive, 53 negative), with the strongest correlations indicating greater abundance in cold-water macroalgae. Salinity had 39 associations (28 negative, 11 positive), likely reflecting the dataset’s predominantly marine sampling (most genomes were from full-salinity ocean environments; freshwater/brackish species had limited representation, see Table S1).

**Figure 2.**
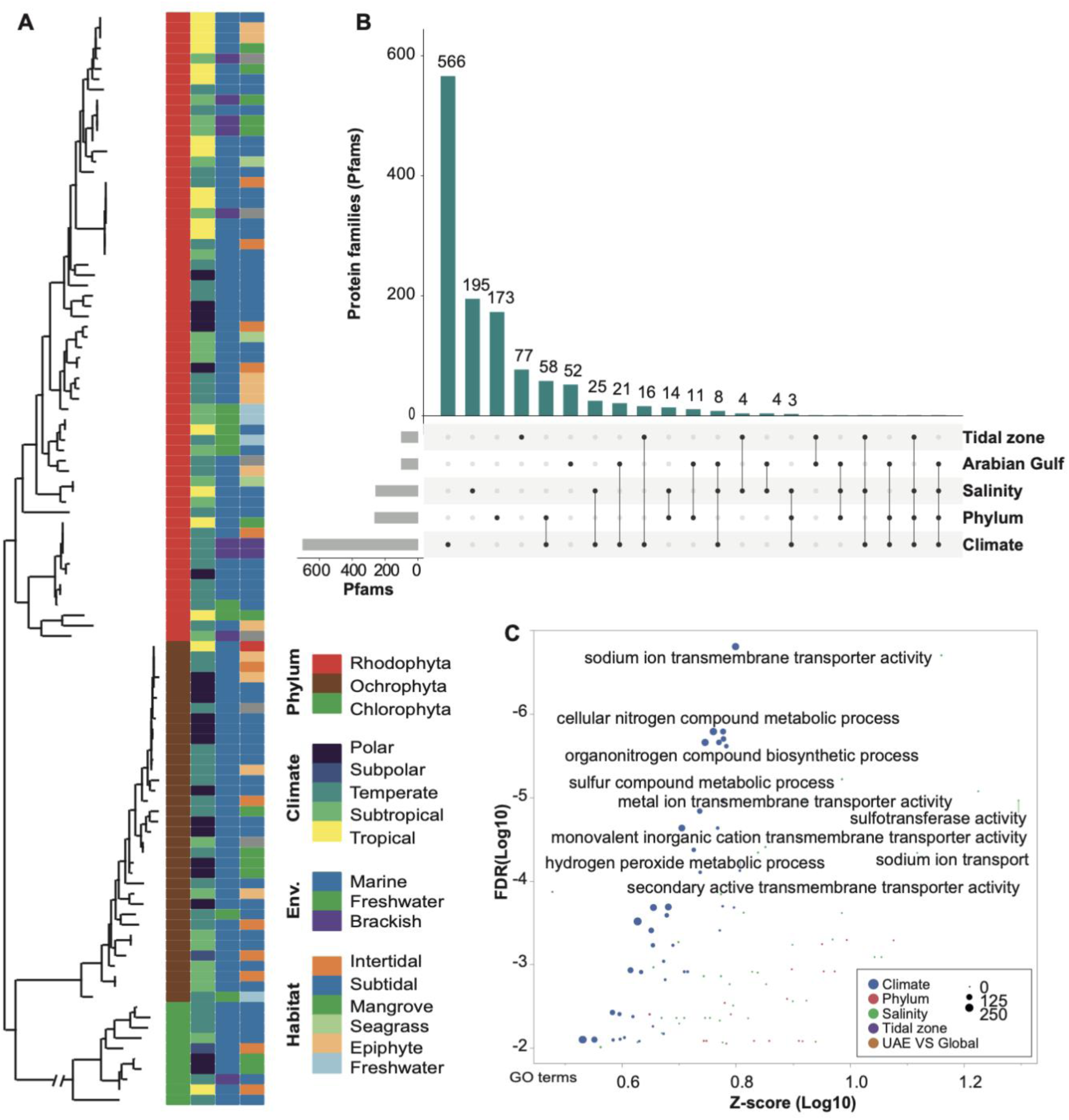
Macroalgal Phylogeny and Pfam Intersect Patterns. (A) Maximum likelihood phylogeny inferred from rbcL protein sequences from the macroalgae used in the study (rbcL markers recovered from 106 of 126 genomes). Node support values represent SH-like local support based on 1000 resamples (values ≥0.80 indicate strong support). Tip colors indicate phylum: Rhodophyta (red, n=61), Ochrophyta (brown, n=35), and Chlorophyta (green, n=10). Adjacent heatmap columns show metadata annotations: climate zone (viridis palette; Polar to Tropical), environment (Marine, Freshwater, Brackish), and habitat type (Intertidal, Subtidal, Mangrove, Seagrass, Epiphyte, Freshwater). (B) UpSet plot showing intersections of significantly enriched PFAM domains across environmental categories. Bar height indicates the number of PFAM families unique to each intersection. Connected dots denote set membership. (C) dcGO enrichment analysis of environment-associated Pfams. Points represent individual GO terms; x-axis shows enrichment Z-score (log10), y-axis shows significance (−log10 FDR). Point size indicates number of associated genes. Colors denote the environmental variable from which PFAM domains were derived: Climate (blue), Phylum (red), Salinity (green), Tidal zone (purple), UAE vs Global (orange).

**Figure 3.**
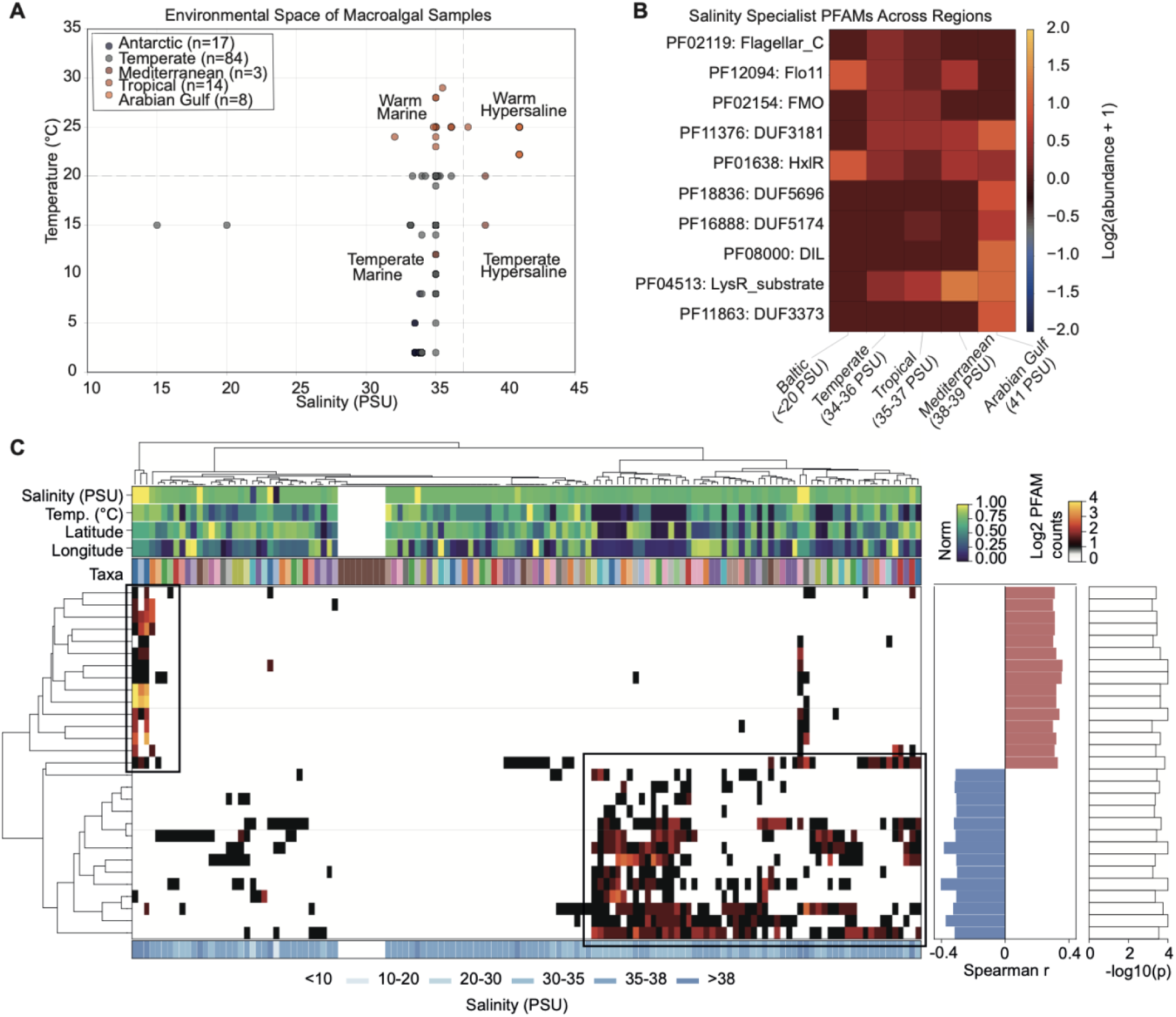
Environmental variable-Pfam untargeted correlation analysis reveals salinity-responsive protein families. A. Environmental space of 126 macroalgal genomes plotted by temperature^79^ and salinity^80^ gradients. Samples cluster into five distinct environmental regimes: Antarctic (n = 17, blue), Temperate Marine (n = 34, light gray), Mediterranean (n = 3, dark gray), Tropical (n = 64, brown), and Arabian Gulf (n = 9, red). The Arabian Gulf samples occupy an extreme environmental niche characterized by both hypersalinity (39-42)^23–27^ and elevated temperatures (22-35°C annual range),^22^ representing the intersection of multiple environmental stressors. B. Heatmap showing differential abundance of salinity-specialist Pfams across geographic regions. Ten protein families with strongest salinity correlations (|r| > 0.31, FDR < 0.05) are displayed as log2-transformed abundance relative to global mean. Putative halosensitive Pfams (top five rows) including cell adhesion domains (PF12094, Flo11), and oxidative stress enzymes (PF02154, FMO) show systematic depletion in hypersaline environments. Putative halophilic Pfams (bottom five rows) comprising membrane-associated domains (PF08000, DIL), transporters (PF04513, LysR_substrate), and uncharacterized osmotic response proteins (DUF families) exhibit reciprocal enrichment. Arabian Gulf populations show the most extreme pattern with 2.0-fold depletion of halosensitive and 1.8-fold enrichment of halophilic domains. C. Hierarchical clustering of 126 macroalgal genomes based on 43 salinity-correlated Pfams indicates convergent osmoadaptive strategies. Environmental metadata tracks (top) show salinity ranges from marine (30-35 PSU, green) to hypersaline (>38 PSU, dark blue). Taxa bar indicates phylogenetic distribution across Rhodophyta (pink), Ochrophyta (brown), and Chlorophyta (green). The main heatmap displays normalized Pfam abundances with two distinct clusters highlighted: halosensitive cluster (blue box, left) showing systematic depletion with increasing salinity, and halophilic cluster (red box, right) with progressive enrichment. Side panels show Spearman correlation coefficients (r) between each Pfam and salinity (blue: negative, red: positive) with significance levels (-log10(p)). The emergent pattern shows phylogenetically independent evolution of osmotic tolerance mechanisms, with 33% of species adopting specialized strategies (clusters marked by boxes) while 67% maintain intermediate profiles, suggesting substantial genomic reorganization is required for extreme salinity adaptation.

The strongest genome-wide significant association with temperature was PF12094 (domain of unknown function, DUF3570; Spearman r = −0.541, p = 6.1×10⁻¹¹, p_Bonferroni = 2.5×10⁻⁶). Additional Bonferroni-significant associations included: salinity with PF15738 and PF06169 (r = −0.518), PF12544 (r = −0.465), PF08972 and PF18405 (r = −0.461), and PF10685 (r = −0.442); temperature with PF13585 (r = −0.452) and PF01638 (r = −0.424); and longitude with PF01062 (r = +0.443).

### Earth-observation–driven environmental correlates of Pfam diversity

To enable scalable characterization of environmental context across sampled genomes, we extracted satellite-derived oceanographic variables using Google Earth Engine (GEE). Based on multiple large-scale, high quality oceanographic datasets, GEE provides global, temporally composited measurements with standardized quality control, enabling direct comparison across 126 macroalgal sampling locations spanning six continents. Multi-year compositing minimized noise from transient oceanographic events while preserving broad environmental gradients relevant to macroalgal ecology.

We evaluated 13 GEE-derived oceanographic variables at 4 km resolution across four-year temporal windows for each location (Table S2), including sea surface temperature metrics (mean, maximum, minimum, annual range, summer, winter), chlorophyll-a concentration (mean, maximum, standard deviation), particulate organic carbon (mean), depth, distance to coast, and water clarity ratio.

Spearman rank correlation analysis between Pfam domain abundances and environmental variables identified 259 significant domain–environment associations (p < 0.001). False discovery rate correction (Benjamini-Hochberg) was applied to the 83,170 tests performed; 12 associations remained significant (q < 0.05), representing 5 unique protein domain families: PF16087 (4 SST associations), PF01638 (4 SST associations), PF14317 (2 SST associations), PF00346 (depth), and PF12094 (SST maximum).

SST-related variables dominated the results (187 of 259 associations at p < 0.001): mean SST had 56 associations, winter SST had 52, and minimum SST had 40. GEE-derived SST correlations were predominantly positive (189 of 259), with the strongest being PF16087 × sst_winter_c (r = +0.582, p = 1.8×10⁻¹⁰). Depth had 32 significant associations. Only two associations survived genome-wide Bonferroni correction (α = 0.05/83,170 = 6.0×10⁻⁷): PF16087 with sst_winter_c (r = +0.582) and sst_max_c (r = +0.467). The PF16087 domain represents beta-loop-helix structures commonly found in transcription factors. Its dominant correlation with GEE variables alludes to the adaptability of eukaryotic genomes enabled through flexible transcriptional programming.

Ignoring phylogenies, the GEE-Pfam matrix correlation analysis revealed a small, statistically robust set of protein domains whose distributions covary with large-scale oceanographic gradients, most strongly with temperature and biogeographic structure. These results highlight a thermally biased signal driven by cold-water lineages, a limited role for salinity, and an unexpectedly high representation of uncharacterized or lineage-specific domains among the top associations. Thus, the satellite-derived environmental framework identifies the principal environmental axes shaping Pfam diversity in macroalgae.

### Per-phylum GEE-based Pfam–environment associations

To distinguish phylogenetically convergent from lineage-specific environmental responses, we performed Pfam–environment correlation analyses stratified by phylum (Rhodophyta, Ochrophyta, Chlorophyta) using Spearman rank correlations, followed by meta-analyses across phyla (Figure 4). Within-phylum correlations used nominal significance (p < 0.05) to preserve sensitivity for the subsequent meta-analysis, which provides multiple testing control through Stouffer’s method. After FDR correction on meta-analysis p-values, 157 associations remained significant (q < 0.05), including 130 convergent associations. The top convergent domains (PF12094, PF01638, PF19572, PF11376) all survived FDR correction (q < 0.01).

**Figure 4.**
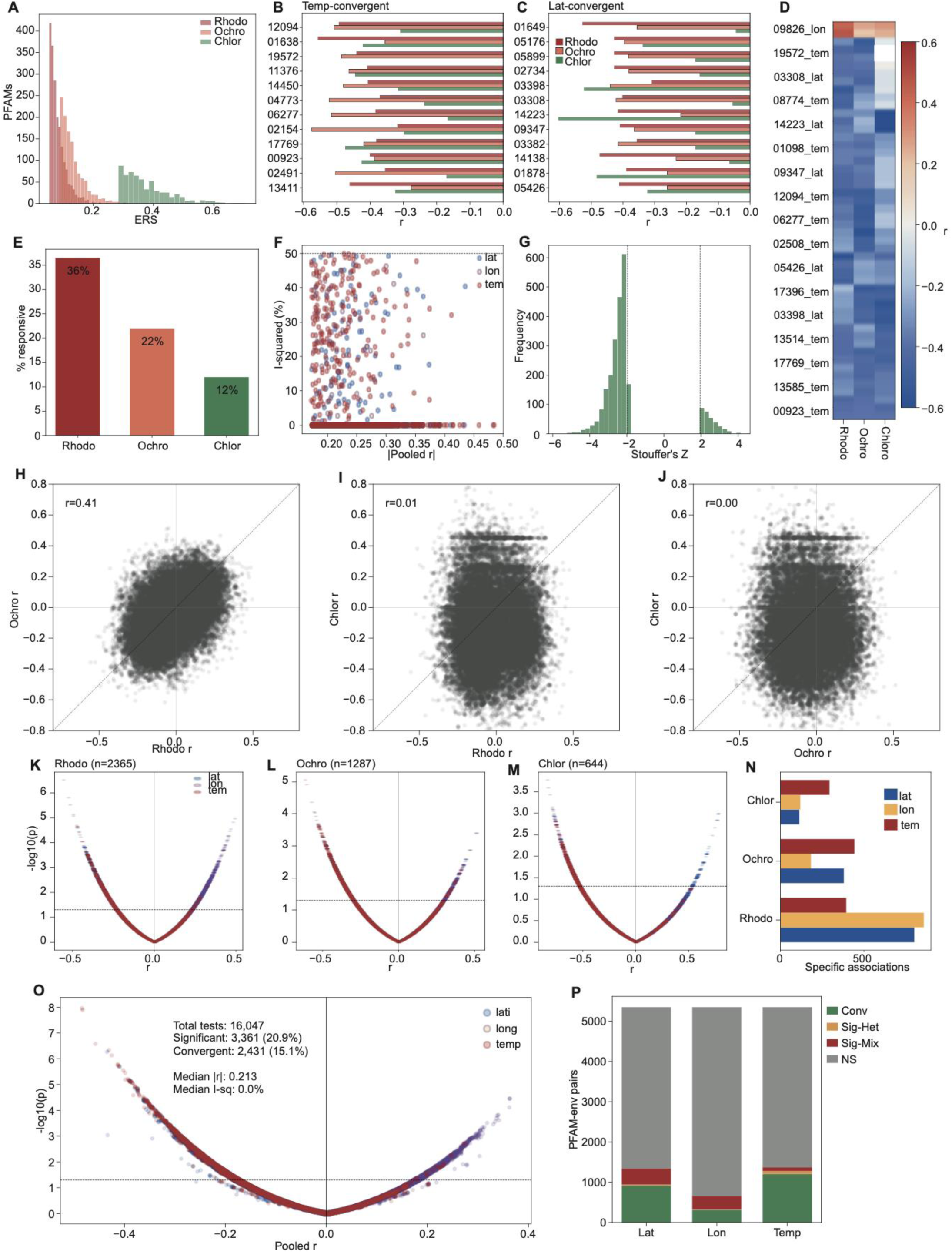
Per-phylum genome–environment (GEE–Pfam) association analyses reveal convergent and lineage-specific patterns. (A) Distribution of environmental response strength (ERS) values for Pfam–environment associations stratified by phylum (Rhodophyta, Ochrophyta, Chlorophyta), summarizing the magnitude of per-Pfam correlation across environmental variables. (B–C) Examples of Pfams exhibiting convergent negative associations across phyla with temperature (B) or latitude (C), with bars indicating per-phylum Spearman correlation coefficients (r). (D) Heatmap of selected Pfam–environment correlations across phyla, illustrating concordant and discordant effect directions. (E) Proportion of Pfams classified as environment-responsive within each phylum. (F) Relationship between pooled absolute correlation strength and variance explained (r²) for latitude, longitude, and temperature. (G) Distribution of Stouffer’s Z scores from per-phylum meta-analysis, with dashed lines indicating significance thresholds. (H–J) Pairwise comparisons of Pfam–environment correlation coefficients between phyla, showing moderate concordance between Rhodophyta and Ochrophyta and weaker correspondence involving Chlorophyta. (K–M) Volcano plots of Pfam–environment correlations for each phylum, showing effect sizes and statistical significance across environmental variables. (N) Counts of phylum-specific Pfam associations by environmental variable. (O) Meta-analysis summary of all Pfam–environment tests, showing pooled effect sizes, significance, and the subset classified as convergent across phyla. (P) Partitioning of Pfam–environment pairs into convergent, heterogeneous, mixed, or non-significant categories for latitude, longitude, and temperature.

Across 16,047 Pfam–environment tests (5,349 per environmental variable), 3,361 (20.9%) had significant associations in the combined meta-analysis (Stouffer’s Z, p < 0.05), with 2,431 (15.1%) classified as convergent based on consistent effect direction across phyla and low heterogeneity (I² < 0.5; Figure 4O–P). Temperature accounted for the majority of convergent associations (1,203), followed by latitude (913) and longitude (315). Convergent associations had a median effect size of |r| = 0.215 and median heterogeneity of I² = 0%, indicating highly consistent effects across lineages (Figure 4G).

The strongest convergent temperature associations were observed for PF12094 (pooled r = −0.484, p < 10⁻⁸), PF01638 (r = −0.483, p < 10⁻⁸), PF19572 (r = −0.458, p < 10⁻⁷), and PF11376 (r = −0.431, p < 10⁻⁶), all showing consistent negative correlations across phyla, indicating enrichment in cold-water lineages (Figure 4B). For latitude, top convergent domains included PF01649 (r = −0.435, p < 10⁻⁶), PF05176 (r = −0.410, p < 10⁻⁵), and PF05899 (r = −0.392, p < 10⁻⁵; Figure 4C). Environmental responsiveness varied substantially across phyla (Figure 4A, E). In Rhodophyta, 36% of Pfam domains (1,965 of 5,388) exhibited at least one significant environmental correlation (p < 0.05), compared to 22% in Ochrophyta (1,159 of 5,285) and 12% in Chlorophyta (609 of 5,076). Pairwise comparison of within-phylum correlation coefficients revealed moderate concordance between Rhodophyta and Ochrophyta (r = 0.41), but weak correspondence involving Chlorophyta (Rhodophyta–Chlorophyta r = 0.01; Ochrophyta–Chlorophyta r = 0.00; Figure 4H–J), suggesting that green algae exhibit distinct environmental response patterns that cannot be captured with their low relative representation in the dataset.

Phylum-specific associations revealed 3,580 Pfam–environment pairs: 2,050 Rhodophyta-specific, 1,005 Ochrophyta-specific, and 525 Chlorophyta-specific (Figure 4N). Among the 3,361 significant meta-analyses results, 814 had significant but mixed-direction effects (opposite correlations in different phyla), and 116 had significant heterogeneous effects (high I² despite consistent direction; Table S2, Figure 4P). These patterns highlight that while convergent environmental tracking predominates, a subset of Pfam domains exhibits lineage-specific or antagonistic responses to environmental gradients.

### The von Willebrand Factor type-A (vWF-A) domain in extreme environment macroalgae

The von Willebrand factor type-A (vWF-A) domain (PF00092) is a conserved protein interaction module that mediates protein–protein and protein–matrix interactions via metal-ion–dependent adhesion sites (MIDAS), providing a plausible molecular basis for enhanced tissue cohesion and substrate attachment under physically and chemically stressful marine conditions.^35–37^

Independent abundance analyses indicated elevated PF00092 copy numbers in Arabian Gulf macroalgae relative to global genomes (2.15-fold; mean 26.6 vs 12.4 copies, n = 9 vs 117; Mann-Whitney U p = 0.038, one-tailed; permutation p = 0.050; bootstrap 95% CI for fold-enrichment [0.96, 3.70]). The bootstrap confidence interval marginally excludes unity at the lower bound, indicating borderline statistical support. Within-phylum comparisons addressed whether PF00092 enrichment reflects phylogenetic composition (UAE samples being enriched for Ochrophyta) or genuine environmental association. UAE Ochrophyta had 3.77-fold higher vWF-A copy numbers than non-UAE Ochrophyta (mean 21.8 vs 5.8 copies; Mann-Whitney p = 0.093, n = 6 vs 38), while UAE Rhodophyta had 2.89-fold enrichment over non-UAE Rhodophyta (mean 49.5 vs 17.1 copies; p = 0.022, n = 2 vs 71). The within-Ochrophyta enrichment exceeded the overall enrichment (3.77× vs 2.15×), and the Rhodophyta pattern reached significance despite only two UAE samples. If the signal were purely phylogenetic, within-phylum comparisons would show no difference. These results suggest environmental rather than phylogenetic drivers, though limited UAE sample sizes preclude definitive conclusions about convergent evolution across all lineages.

Across all samples, PF00092 showed biogeographic associations with latitude (r = −0.295, p = 0.0008, n = 126) that reached nominal significance but did not survive FDR correction. Environmental Responsiveness Scores (ERS; mean R² of significant correlations) revealed phylum-specific patterns: Rhodophyta showed moderate responsiveness (ERS = 0.106) with geographic associations (latitude r = −0.325, longitude r = +0.325; both p = 0.005), while Chlorophyta and Ochrophyta showed no significant environmental correlations (ERS = 0.0). Within Ochrophyta specifically, latitude association was stronger (r = −0.334, p = 0.0003) with weaker temperature correlation (r = +0.250, p = 0.008).

Direct temperature correlations were weak across all phyla (max |r| = 0.23), suggesting the Arabian Gulf enrichment reflects regional factors rather than simple thermal gradients. Moran’s I analysis confirmed no significant spatial clustering (I = 0.029, p = 0.44), indicating this pattern is not an artifact of geographic autocorrelation.

Although vWF-A domains are found across Eukaryota, Eubacteria, and Archaea, their functional roles in macroalgae remain uncharacterized.^3,10,19,35–39^ The conserved MIDAS motif within vWF-A domains mediates divalent cation-dependent adhesion to extracellular matrix proteins and other ligands, supporting cell–cell cohesion and substrate attachment across diverse taxa.^40–42^

The Arabian Gulf’s shallow coastal waters (< 30 m depth at all sampling sites) experience substantial hydrodynamic stress from frequent shamal wind events that generate strong wave action and recurrent sediment resuspension.^28^ Environmental extremes characteristic of this region, including summer temperatures exceeding 35 °C and salinities above 44 PSU,^28^ may further compound mechanical stress by altering cell junction integrity, tissue hydration, and adhesion kinetics. Enhanced vWF-A copy numbers may therefore reflect selection for reinforced substrate attachment under chronic mechanical forcing combined with thermal and osmotic stress—a hypothesis consistent with the domain’s well-established adhesive function.

### AlphaEarth vision transformer environmental embedding correlations

To discover additional environment-genomic^43^ relationships, we applied AlphaEarth Foundations (AEF),^44^ a vision transformer (ViT) model^45^ trained on petabyte-scale satellite imagery datasets,^46^ to extract 64-dimensional environmental embeddings at 10m resolution for each sampling location (2017-2024 composite; see Tables S5-S9). Vision transformer embeddings are learned representations beyond remote sensing measurements of specific physical variables (e.g., sea surface temperature in °C) that encode complex, multivariate patterns visible in satellite imagery. This approach enabled discovery of genome-environment associations beyond those detectable with traditional single-variable analyses (Table 1).

**Table 1.** Environmental correlations and ecological interpretations of top AEF embedding dimensions. Spearman rank correlations between AEF embedding dimensions and environmental variables (Temperature, Latitude, Longitude) across 126 macroalgal samples (70 Rhodophyta, 43 Ochrophyta, 13 Chlorophyta). Dimensions ranked by absolute correlation strength with their dominant environmental axis. “Strongest Variable” indicates the environmental variable with the highest |r| for each dimension. PFAM associations represent the number of protein domains significantly correlated with each dimension (FDR < 0.05; 313 total associations across all dimensions). Ecological interpretations derived from correlation sign and magnitude: negative temperature correlations indicate cold-water enrichment; negative latitude correlations indicate high-latitude/polar enrichment; positive values indicate the inverse.

| AEF DIM | STRONGEST VARIABLE | TEMPERATURE R | LATITUDE R | LONGITUDE R | PFAMS | ECOLOGICAL MEANING |
| --- | --- | --- | --- | --- | --- | --- |
| A08 | Temperature | -0.685 | 0.074 | -0.364 | 6 | Cold-water indicator |
| A14 | Latitude | 0.031 | -0.59 | 0.116 | 4 | High-latitude / polar gradient |
| A52 | Temperature | -0.703 | 0.17 | -0.305 | 8 | Cold-water indicator |
| A05 | Latitude | -0.098 | -0.555 | -0.059 | 0 | High-latitude / polar gradient |
| A26 | Latitude | 0.059 | 0.552 | 0.304 | 0 | Low-latitude / tropical gradient |
| A41 | Temperature | -0.544 | 0.262 | -0.306 | 1 | Cold-water indicator |
| A49 | Latitude | 0.13 | -0.536 | 0.161 | 0 | High-latitude / polar gradient |
| A00 | Temperature | 0.511 | -0.074 | 0.339 | 2 | Warm-water / subtropical indicator |
| A10 | Latitude | 0.081 | -0.504 | 0.028 | 1 | High-latitude / polar gradient |
| A53 | Temperature | 0.49 | -0.314 | 0.134 | 31 | Warm-water gradient |
| A28 | Latitude | 0.032 | 0.488 | 0.047 | 0 | Subtropical-temperate transition |
| A48 | Latitude | 0.138 | 0.476 | -0.154 | 4 | Subtropical-temperate transition |

All 64 AEF dimensions (100%) correlated significantly (p < 0.05) with at least one GEE environmental variable (Table S3), with 43 dimensions (67.2%) surviving Bonferroni correction (p < 6.0×10⁻⁵). Sea surface temperature correlated with 58 dimensions (strongest: A52, r = -0.703, p < 10⁻^15^), distance from coast with 42 dimensions (strongest: A14, r = 0.600), depth with 39 dimensions (strongest: A14, r = 0.585), chlorophyll concentration with 38 dimensions (strongest: A54, r = 0.580), and particulate organic carbon with 30 dimensions (strongest: A03, r = 0.659). The remaining 12 dimensions (A11, A19, A20, A31, A34, A37, A38, A40, A55, A57, A58, A59) had no significant correlation with mean temperature, latitude, or longitude, but correlated with derived GEE oceanographic features not represented in simple collection metadata. Specifically: A11 correlated with SST annual range (r = -0.33, p = 0.002), capturing thermal seasonality rather than mean temperature; A20 correlated strongly with ocean productivity variables including chlorophyll concentration (r = -0.56), particulate organic carbon (r = -0.55), and water clarity (r = 0.58); A31 correlated with depth (r = 0.44) and distance to coast (r = 0.48), representing a nearshore-to-offshore gradient; A57 captured thermal variability through correlations with SST minimum, annual range, and winter temperatures; and A19, A34, A55, A58, and A59 correlated primarily with summer SST and coastal proximity. These patterns indicate that AEF dimensions partition environmental space along axes orthogonal to simple latitudinal or mean-temperature gradients, including seasonal thermal amplitude, productivity regimes, and coastal-oceanic transitions (Table S3).

Correlations between AEF dimensions and GEE-derived oceanographic variables (e.g., A52 with SST, r = -0.703; Table S3) show that the vision transformer’s learned representations encode biologically-relevant environmental variation rather than image artifacts. Traditional correlation analysis using manually-collected environmental metadata (collection temperature, latitude, longitude) identified fewer Pfam associations and failed to detect the 12 AEF dimensions (A11, A19, A20, A31, A34, A37, A38, A40, A55, A57, A58, A59) that capture complex gradients such as seasonal SST variability, coastal proximity, bathymetry, and productivity—environmental features not represented in simple collection metadata but critical for understanding genomic adaptation.

Within-phylum analysis of Ochrophyta revealed only two Pfam domains surviving FDR correction (q < 0.05), both strongly associated with dimension A56: NAD kinase (PF01513; r = −0.703) and Drought-induced 19 protein (Di19; PF05605; r = −0.702). These domains clustered together in bi-cluster module 9, indicating coordinated environmental response. NAD kinase catalyzes the phosphorylation of NAD to NADP, a rate-limiting step in NADPH production essential for both photosynthetic electron transport and ROS detoxification under stress. Di19 encodes a zinc-finger transcription factor first characterized in drought-stressed *Arabidopsis*, where it regulates osmotic and oxidative stress responses. The co-occurrence of these two domains—one metabolic, one regulatory—suggests a coordinated stress-response module in brown algae linking energy metabolism (NADPH supply) with transcriptional regulation of osmotic tolerance. Dimension A56 correlates with coastal proximity and productivity gradients in GEE, consistent with intertidal brown algae experiencing cyclical desiccation and osmotic stress during tidal exposure.

Restricting analysis to well-represented PFAMs (≥10 non-zero samples) revealed substantially more significant associations within Rhodophyta: 1,036 significant PFAM-dimension correlations (FDR<0.05) compared to 137 under the permissive filter used for cross-phylum comparison. This difference reflects both improved correlation reliability for well-represented domains and reduced multiple testing burden.

### Pfam matrix–AEF associations reveal climate-driven Pfam modules

Several protein families had significant correlations with AEF dimensions (Table S3). The AhpC/TSA peroxidase domain (PF13411) increases along dimension A18 (r = 0.479, p = 1.4×10⁻⁸), which correlates with decreasing latitude (r = -0.352), indicating enrichment in tropical waters. Conversely, the helix-turn-helix domain (PF01638) decreases in warm, saline conditions (r = -0.472 with dimension A53, which correlates with temperature r = 0.394 and salinity r = 0.297), suggesting specialization to cooler, fresher coastal waters. Notably, the NAD(P)-binding domain (PF10988) correlates with dimension A36 (r = 0.463, p = 4.9×10⁻⁸), a dimension uncorrelated with measured variables—this association may reflect adaptation to ocean productivity gradients visible as color in satellite imagery but absent from *in situ* measurements.

The correlation analysis produced a structured matrix linking macroalgal genomic composition to latent environmental features derived from AEF embeddings. Most Pfam–dimension pairs had weak or near-zero correlations, likely reflecting core housekeeping functions independent of habitat. A minority displayed strong positive or negative associations (|ρ| > 0.25, FDR < 0.05), signifying functional domains that covary with environmental gradients. Examples include a Pfam exhibiting strong positive correlation with tropical-associated dimension A06 (ρ = +0.30, p < 0.001) suggesting gene family expansion in low-latitude species, and negative correlation with polar-associated dimension A01 (ρ = −0.41, p < 0.001) implying selective loss in cold environments. When visualized after z-score normalization, these associations formed coherent bi-clusters corresponding to gene-environment modules.

Bi-clustering^47,48^ of the AEF–Pfam correlation matrix revealed two major blocks of coordinated genomic–environmental associations (Fig. 5). The first cluster (blue, “tropical-depletion”) spans AEF dimensions A06, A07, A12, A18, and A53, all showing strong negative correlations with latitude (r = – 0.24 to –0.40, p < 0.05–0.001). A total of 118 unique Pfams had significant correlations (FDR < 0.05) with at least one of these dimensions (128 total associations), with A18 alone explaining 48 associations. Pfams in this cluster display negative z-scores, indicating functional depletion in warm-water environments. The second cluster (tan–olive, “polar-enrichment”) spans dimensions A17, A25, A30, and A46, each exhibiting positive correlations with latitude (r = +0.24 to +0.33, p < 0.05–0.001). This cluster comprises 67 unique Pfams (79 total associations) with A25 contributing 49 associations—the highest of any AEF dimension (Table S3). Pfams within this module exhibit positive z-scores, reflecting systematic enrichment in cold-water macroalgae. The emergence of discrete, non-random bi-clusters demonstrates that unsupervised environmental embeddings derived from satellite data predict functional genomic composition across marine macroalgae.

**Figure 5.**
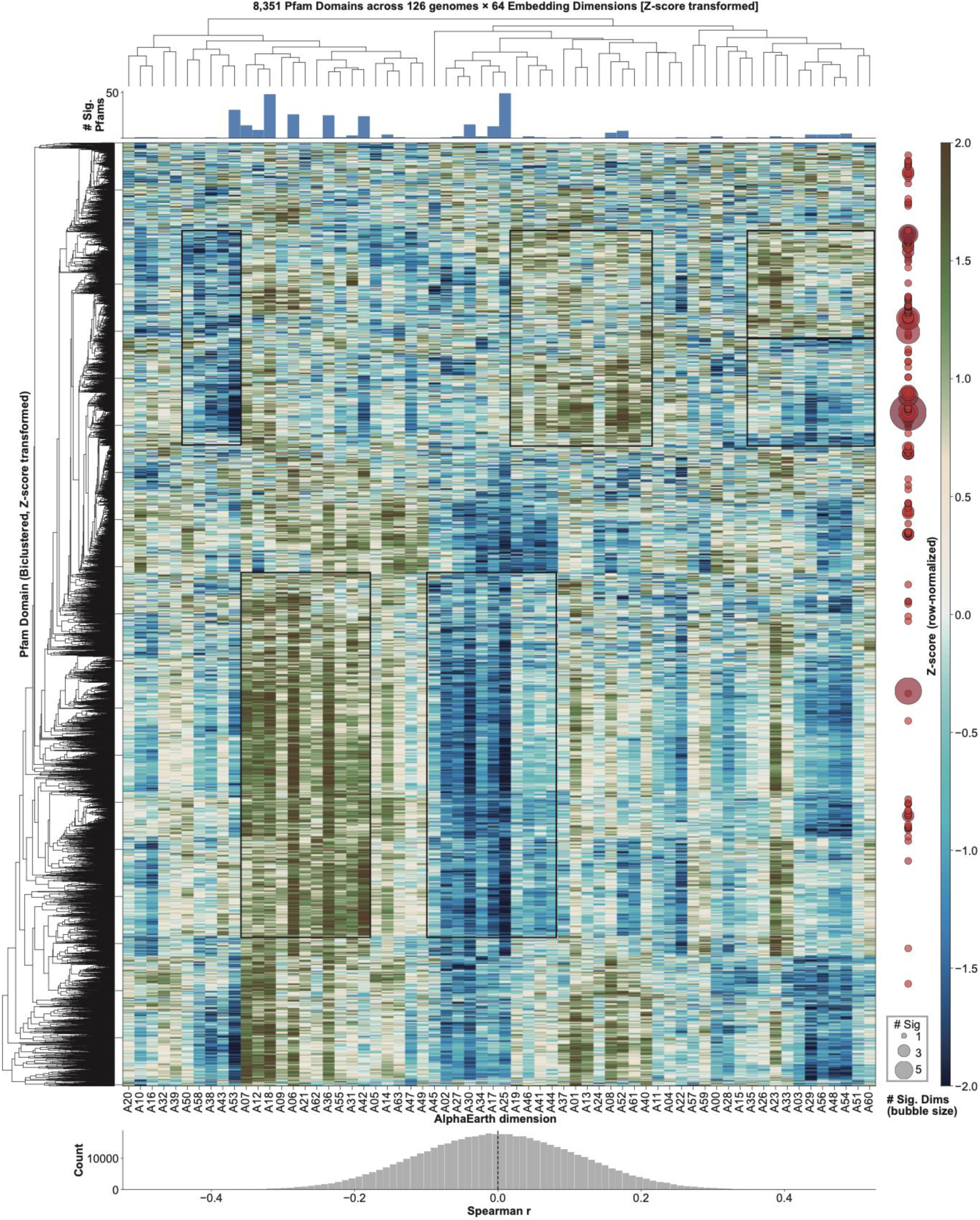
AEF dimension correlation landscape reveals Pfam modules responsive to multidimensional environmental gradients. Hierarchically bi-clustered heatmap showing Spearman rank correlations between 10,300 Pfam protein domain families and 64 AEF vision transformer embedding dimensions across 126 macroalgal genomes. AEF dimensions (A00-A63, x-axis) represent learned environmental representations extracted from 10-meter resolution Sentinel-2 satellite imagery at genome collection sites via a pre-trained vision transformer model. Pfam domains (y-axis) quantify functional genomic content, with abundance values normalized as counts per genome. Both axes are ordered by hierarchical clustering (Ward linkage, Euclidean distance on z-score transformed correlation values) to reveal co-varying functional modules. Color Scale: Heatmap cells represent Spearman’s rank correlation coefficient (ρ) ranging from -0.8 (strong negative correlation, ocean blue) to +0.8 (strong positive correlation, earth brown), with white indicating no correlation (ρ = 0). Each cell represents one of 659,200 Pfam-dimension correlation tests (10,300 domains × 64 dimensions). Spearman correlation was used to accommodate non-normal distributions and outliers in domain count data. False discovery rate (FDR) correction was applied using the Benjamini-Hochberg method^78^ across all tests.

### Gene ontology overlap between saltwater species-specific Pfams and AEF dimension A06

We used Pfams unique to macroalgae from each environmental category (Fig. 2) and performed intersect analyses with Pfams significantly correlating with AEF embedding dimensions. The strongest overlap signal was from dimension A06-significantly correlated Pfams and those unique to saltwater species (Figs. 2,5). We then performed dcGO enrichment ^49^ analyses on these Pfams (Table S3). We identified GO terms from Pfams that are both unique to saltwater macroalgae and significantly correlated with the AEF A06 embedding dimension. These terms are dominated by ion transport functions (13/17 terms, 76%), particularly cation transmembrane transport, coupled with energy metabolism through ATP synthase complexes. Four core Pfam domains (PF00006, PF02823, PF02874, PF00230), representing ATP synthase subunits and aquaporin-like channels, appear consistently across these terms, suggesting a convergent molecular strategy for osmotic homeostasis in marine environments.

## DISCUSSION

### Integrating functional genomics and remote sensing

Because satellite products—such as temperature, salinity, irradiance, turbidity, and nutrient proxies—capture multi-year, landscape-scale environmental pressures, they reflect selective forces shaping genome content over evolutionary time. Traditional phylogenetic analyses recover signals tied to ancestry rather than adaptive function—and then only in examined genes and not in a genome-wide context. In contrast, Pfam–environment correlations highlight functionally equivalent molecular strategies that appear independently in distantly related taxa exposed to similar conditions, such as high-salinity stress proteins in both Chlorophytes and Rhodophytes inhabiting hypersaline niches.

This approach reveals convergent solutions rather than lineage history, enabling detection of adaptive strategies that transcend phylogenetic boundaries.

Many environmental adaptations involve coordinated shifts across dozens of protein families with small effects. Marker-gene analyses—such as those based on rbcL or 18S rRNA—cannot capture these distributed signatures, while Pfam patterns integrate signals from many loci, increasing power to detect subtle, polygenic adaptations. Furthermore, satellite data provides high-resolution, quantitative gradients in variables like temperature variance, photosynthetically active radiation, chlorophyll concentration, and wave energy. These continuous measurements enable detection of dose–response-like genomic patterns that reveal thresholds or smooth clines in domain prevalence, reflecting long-term environmental optimization.

Importantly, genomes encode traits tuned to the actual conditions organisms experience, and satellite products measure these conditions directly—often over decades—capturing the persistent, realized environmental envelope better than point-sampling or laboratory measurements. Long-term irradiance variability, seasonal turbidity regimes, episodic heat waves, or chronic nutrient stress can leave distinct genomic fingerprints not evident in snapshot environmental metadata or conventional ecological categories. These selective pressures, invisible to classical niche descriptors, become detectable through the integration of functional genomics and remote sensing.

Because Pfam domains represent functional units, they allow alignment of adaptive signals across Rhodophytes, Chlorophytes, and Phaeophytes—something traditional marker genes cannot resolve without phylogenetic correction. Convergent enrichment patterns across clades indicate true environmental adaptation rather than shared evolutionary history. In essence, by fusing functional genomic descriptors with multi-year satellite observations, we can uncover deep, convergent genomic signatures associated with environmental pressures that remain invisible to phylogenies, short-term sampling, or single-gene markers.

### Methodological relationship between GEE and AEF frameworks

The strong correlation between AEF dimensions and GEE-derived variables, such as the correlation of negative 0.703 between A52 and Sea Surface Temperature, confirms that vision transformer embeddings capture tangible oceanographic signals rather than stochastic image artifacts. This alignment establishes a convergent validation strategy where the concordant Pfam associations across both frameworks demonstrate robust analytical consistency. The dual-framework approach provides a multi-scale perspective because AEF provides a high-resolution view at 10 meters that captures habitat heterogeneity lost in the 4-kilometer products of GEE. Furthermore, AEF dimensions integrate nonlinear, high-dimensional patterns across spectral bands to identify complex environmental signatures that often escape standard single-variable parameterization. Associations unique to the AEF framework highlight adaptive responses to visual features that are not captured by traditional sensors, while GEE provides the physically-grounded interpretive anchor. Ultimately, AEF extends the reach of the analysis to patterns that resist explicit parameterization, and the convergence of these frameworks increases the statistical confidence that the identified genomic associations are fundamental to the biology of the organism rather than artifacts of a specific analytical pipeline.

### Implications for climate change predictions and biotechnology

The modular genomic architecture of environment-protein domain associations enables multidimensional environmental tolerance screening.^50^ A species with strong representation of thermal-associated domains (high PF06415 iPGM, PF01638 HxlR) but weak representation of osmotic-associated domains (low PF00330 Aconitase, PF02347 Glycine cleavage) might tolerate temperature changes but face challenges under combined ocean warming and altered salinity patterns (from increased precipitation) or ocean acidification (which can generate secondary osmotic stress through ion dysregulation).^51^

The identification of distinct genomic signatures for environmental associations, validated across both traditional oceanographic variables and machine learning-derived environmental representations, provides a molecular framework for investigating macroalgal genomic composition across environmental gradients.^52^ The modular nature of genomic variation we discovered suggests that species responses to environmental change may depend on their existing genomic composition.^53^

This work demonstrates that macroalgae from extreme environments harbor genomic features detectable through both interpretable (GEE) and learned (AEF) environmental representations. As we continue to face environmental growing change, preserving these genomic resources becomes imperative.^54^ The Arabian Gulf’s macroalgae today inhabit environmental conditions that are representative of future ocean states in some regions^55^ and possess genes potentially relevant to stress tolerance. Their conservation is both an ecological and practical consideration for maintaining options for ecosystem restoration and sustainable aquaculture.^56^ The consistency of findings across independent analytical approaches—traditional oceanography and machine learning—strengthens confidence in these environment-genomic associations.

### Limitations of the study

While this work establishes a framework for macroalgal genomic ecology, the scale of current sampling (n=126) and the dominance of specific phyla provide a foundational, rather than exhaustive, characterization of these lineages. Our findings reflect the visual and environmental signals captured at the point of sampling; however, we recognize that the complex interplay of evolutionary forces—from genetic drift to lineage sorting—underlies the observed Pfam variation. By controlling for phylogenetic hierarchy and spatial autocorrelation, we provide a conservative baseline for genome-environment associations. These results offer a predictive roadmap for future high-density sampling efforts aimed at isolating direct adaptive mechanisms from broader evolutionary noise.

## METHODS

### Sample collection and species identification

To supplement the global macroalgal genome dataset with lineages from hypersaline subtropical habitats, we sampled along the United Arab Emirates (UAE) coast of the Arabian Gulf. Collections were performed during winter months (water temperatures 22–25°C), representing the season of peak macroalgal diversity and abundance in this region. We collected nine target species from four locations—Al Hiel, Dhabiya, Ras Ghurab, and North Corniche Abu Dhabi—spanning intertidal and subtidal zones to depths of seven meters (Figure 1A, Table S1).

Specimens were identified via morphological and molecular assessment, revealing a diverse assemblage spanning all three major macroalgal phyla: six brown algae (*Padina boergesenii*, *Polycladia myrica, Sargassum latifolium, S. angustifolium, Canistrocarpus cervicornis,* and one unidentified endophyte), one red alga (*Chondria dasyphylla*), and one green alga (*Avrainvillea amadelpha*). These taxa have been previously documented in UAE and Iranian waters,^57,58^ confirming established adaptation to Gulf conditions. Notably, *Padina* and *Sargassum* specimens were sampled from degraded substrates on recently bleached coral reefs, while *A. amadelpha* was identified consistent with invasive lineages described in the Mediterranean and Hawaii.^34^ These genomes were treated as public data in prior comparative analyses; however, this study represents their first primary description.^32^

### DNA extraction and sequencing

To mitigate contamination from natural symbioses (bacteria, fungi, endophytes) and extraction inhibitors (polysaccharides, polyphenols), we selected the least-affected tissues and treated samples with 0.1% SDS prior to lysis. Genomic DNA was extracted using a modified CTAB-based phenol–chloroform protocol. Whole-genome sequencing libraries were prepared and sequenced at New York University Abu Dhabi (NYUAD) on an Illumina NovaSeq PE150 platform, targeting approximately 100× coverage per genome (NCBI BioProject: PRJNA929663; Table S1).

### Assembler benchmarking and genome assembly

To select an optimal assembly strategy, we benchmarked three assemblers—CLC Genomics Workbench, ABySS, and SPAdes—using raw reads from eight representative macroalgae and *Chlamydomonas reinhardtii (*Table 1). ABySS and SPAdes were executed on HPC systems, while CLC was run locally. We evaluated performance based on N50 values and BUSCO completeness (Eukaryota dataset with AUGUSTUS) across k-mers 40–60 (Table S1, Fig. S1). While SPAdes achieved high completeness for specific small genomes, it failed on larger genomes. CLC consistently generated higher N50 values and comparable or superior BUSCO scores for most red, green, and brown macroalgae (Table S1; Figure S1).

Based on these results, the final nine UAE genomes were assembled using CLC Genomics Workbench. Reads were trimmed to remove adapters, terminal bases, homopolymers, and polynucleotide runs. Assemblies were performed using a k-mer sweep (40, 50, 60) with a minimum contig length of 500 bp and paired-distance autodetection enabled. To remove non-target organism contigs, all assemblies underwent six rounds of decontamination using the BLAST-Limiting Eukaryotic Assembly Contamination, Heuristically (BLEACH) pipeline.^32^

### Dataset compilation and quality assessment

The nine newly described UAE genomes were integrated with 117 publicly available macroalgal genomes. The mean BUSCO completeness score for the final combined dataset was 68.8 ±15.0 %present (Fig S1). We retained all genomes for downstream analysis regardless of BUSCO score. This decision was based on the observation that standard BUSCO ortholog sets are biased toward model eukaryotes and routinely underestimate completeness in macroalgal lineages.^59,60^ Empirical studies indicate that “incomplete” algal assemblies often retain the majority of metabolic pathways and regulatory modules, and that strictly filtering by completeness results in a greater loss of phylogenetic signal than the inclusion of missing data.^61^ Retaining these genomes ensures that rare and underrepresented clades contribute to comparative analyses.

The reference genomes in this study included: *Ectocarpus siliculosus* (Phaeophyte, ∼ 196 megabase pairs (Mbps), ^10^; *Ectocarpus subulatus* (Phaeophyte, 242 Mbps), ^11^; *Saccharina japonica* (Phaeophyte, ∼ 535 Mbps), ^62^; *Chondrus crispus* (Rhodophyte, ∼ 104.98 Mbps), ^12^; *Gracilariopsis chorda* (Rhodophyte, 92.18 Mbps), ^13^; *Porphyra umbilicalis* (Rhodophyte, ∼ 87.89 Mbps), ^14^; Neopyropia yezoensis (Rhodophyte, ∼107.59 Mb), ^15^; *Neoporphyra haitanensis* (Rhodophyte, 53.25 Mbps), *Gracilariopsis lemaneiformis* (Rhodophyte, 88.69 Mbps), ^16^ and *Kappaphycus alvarezii* (Rhodophyte, 336.72 Mbps), ^17^; *Caulerpa lentillifera* (Chlorophyta, 28.77 Mbps), ^18^; *Ulva mutabilis* (Chlorophyte, 98.48 Mbps), ^19^; *Ulva prolifera* (Chlorophyte, 887.88 Mbps), ^63^; and *Chara braunii* (Chlorophyte/Streptophyte 1751.21 Mbps), ^64^. See Table S1D for more information on the reference genomes used in this study.

### Phylogenetic analysis

To establish the taxonomic categorization of the dataset and verify species identities, ribulose-1,5-bisphosphate carboxylase/oxygenase (RuBisCo) large subunit (rbcL) markers were identified in 116 macroalgal genomes via tBLASTn searches using phylum-specific reference queries: Ectocarpus sp. (AAR24444.1) for Ochrophyta, *Gracilariopsis hommersandii* (AAL46528.1) for Rhodophyta, and Ulva rigida (QWL15241.1) for Chlorophyta.

Identified sequences were aligned using MUSCLE v3.8.31 with default parameters (Data S1). Maximum likelihood phylogenetic reconstruction was performed using Molecular Evolutionary Genetics Analysis (MEGA) v11 with the Jones-Taylor-Thornton (JTT) substitution model and gamma-distributed rates across four categories. The heuristic search utilized the Nearest-Neighbor-Interchange (NNI) method, and nodal support was assessed via 1,000 bootstrap replicates. Final trees were visualized and annotated using the Interactive Tree Of Life (iTOL) v6.

Analyses are presented as ‘pan-macroalgal’ [all samples pooled] and phylum-stratified to address two different scopes of inquiry. Overall, the dataset encompasses 126 macroalgal genomes across the three phyla, so phylum-level splits represent the maximum possible phylogenetic exclusion with present taxon coverage.

### Protein family annotation and analysis

HMMsearch results from Nelson, et al., 2019^32^ were used as inputs for comparative analyses. Pfam domain counts were aggregated at the genome level. This initial matrix contained 10,589 unique Pfam domains (10,300 after filtering to remove all-zero and invariant domains), reflecting the diverse functional repertoire across red, brown, and green macroalgae. The dataset included a composition of macroalgal genomes with unequal phylum representation (Table S1). This imbalance reflects both biological reality—red algae dominate marine macroalgal diversity—and current sequencing priorities. To prevent spurious associations driven by majority-class patterns, we implemented multiple safeguards: (phylum-stratified correlation analyses with sample-size–weighted meta-analysis (Stouffer’s method);^65^ heterogeneity testing (I² statistic)^66^ to reject associations inconsistent across lineages; and within-phylum analyses to confirm lineage-specific signals (Table S3). These approaches ensure that reported associations reflect genuine environment-genome relationships detectable across evolutionary divergences rather than artifacts of sampling bias.

Variance analysis revealed substantial heterogeneity among Pfam domains, with coefficients of variation (CV = σ/μ) ranging from 0.12 to 8.94 (median = 1.23), indicating moderate to high variability across genomes (Table S2). This variability is critical for predictive modeling, as invariant features lack discriminatory power. All data and parameters supporting reproducibility are provided as Supplementary Data files, including the complete Pfam count matrix (126 genomes × 10,589 domains) and the metadata integration table (Table S1).

### Intersection analysis (UpsetR)

All Pfam counts were converted into binary values (0 and 1) with all counts greater than 0 being replaced by 1 for exclusion/UpSetR^67^ analysis. For the tidal zone (intertidal and subtidal), the 18 genomes that were epiphytes (habitat - Table S1) or belong to freshwater environment (salinity - Table S1) were excluded by assigning a value of 0 for all 100 significant Pfams. Then, for each metadata variable (climate, phylum, environment, location, salinity, and tidal zone), we extracted the binary Pfam counts for the above mentioned significant ranked Pfams and saved them as .txt files. The R function ‘upset’ in the UpSetR package was used to generate the UpSetR plot. For the summarized Pfams UpSetR plot (Figure 2A), we extracted the 1266 unique Pfams, then for each Pfam, we assigned 1 or 0 when the Pfam is significant or not for the metadata, respectively. The R function upset was used to generate the UpSetR plots.

### Google Earth Engine oceanography correlations

Google Earth Engine (GEE) is a cloud-based planetary-scale platform for geospatial analysis that provides access to petabyte-scale satellite remote sensing datasets with integrated computational infrastructure.^68^ The platform architecture combines a multi-petabyte data catalog of publicly available satellite imagery and geophysical datasets, distributed computing infrastructure built on Google’s data centers enabling parallel processing across thousands of machines, and JavaScript and Python APIs for specifying geospatial analyses that execute server-side without requiring local data download or storage. The GEE platform integrates multi-decadal satellite time series (MODIS Aqua, Landsat, Sentinel) to quantify sea surface temperature (mean, maximum, minimum, annual range, summer/winter averages), ocean productivity (chlorophyll-a concentration, particulate organic carbon), physical parameters (bathymetry, distance to coast, water clarity), and geographic coordinates (latitude, longitude) at 4km spatial resolution averaged from 2020-2023.

Environmental variables come from four primary satellite products, selected for their relevance to macroalgal ecophysiology, spatial coverage, and temporal availability: Sea Surface Temperature (SST): NASA MODIS Aqua Level-3 Standard Mapped Images (product ID: NASA_OCEANDATA/MODIS-Aqua/L3SMI) provide daily global SST at 4km nominal resolution using the multi-channel SST algorithm.^69^ This algorithm combines thermal infrared measurements at 11μm (band 31) and 12μm (band 32) with atmospheric correction models to retrieve sea surface skin temperature with ±0.5°C accuracy against in situ buoy measurements. We computed four-year temporal composites (2020-2023) yielding mean, minimum, maximum, and standard deviation statistics to capture both average thermal conditions and thermal variability (seasonality, interannual variation). The 4-year window was selected to match typical macroalgal generation times and provide robust estimates resilient to individual extreme events.

Chlorophyll-a and ocean Color Variables: MODIS Aqua Ocean Color products (same collection) provide 8-day composite chlorophyll-a concentrations using the OC3M algorithm,^70^ which relates blue-to-green band ratios (443nm, 488nm, 551nm reflectance) to chlorophyll concentration via empirical relationships calibrated against global in situ datasets. Accuracy is ∼35% median error in open ocean (Case I waters) but degrades to ∼50% in optically complex coastal waters (Case II) with high sediment or colored dissolved organic matter. We also extracted particulate organic carbon (POC) concentrations using empirical carbon algorithms^71^ and photosynthetically available radiation (PAR) from the same product suite, providing integrated measures of primary productivity and light availability.

For bathymetry, NOAA ETOPO1 Global Relief Model provided seafloor depth at 1 arc-minute (∼2km) resolution by integrating ship-based soundings, satellite altimetry, and coastal digital elevation models.^72^ Unlike dynamic oceanographic variables, bathymetry is a static layer representing long-term geomorphology. Depth accuracy varies from ∼100m in deep ocean to ∼10m in well-surveyed coastal regions. Bathymetry serves as a proxy for multiple environmental factors: light attenuation (deeper = lower irradiance), wave exposure (shallow coastal vs. deep offshore), and nutrient dynamics (upwelling zones). Genome–environment association analyses were restricted to 101–105 genomes with complete satellite coverage, as some coastal locations lacked full GEE data.

To focus statistical power on informative associations, we restricted correlation testing to Pfam domains present in ≥5% of genomes (n ≥ 6 samples), excluding rare domains that lack sufficient variance for reliable correlation estimation. This filtering reduced the test space from ∼141,000 potential correlations to 83,170 Pfam–environment pairs actually tested.

### Phylogenetic Correction of Pfam–Environment Associations

To distinguish lineage-independent environmental adaptation from phylogenetic signal, Pfam–environment correlations were evaluated using a phylum-stratified meta-analytic framework (Table S3). For each Pfam–environment pair, Spearman correlations were computed within each phylum independently (Rhodophyta, Ochrophyta, Chlorophyta). This removes between-phylum covariance arising from shared ancestry and biogeographic structure. Correlation coefficients were Fisher Z-transformed and combined across phyla using Stouffer’s Z-score method,^65^ weighting by the square root of sample size. We quantified effect-size consistency using the I² statistic.^66^

The I² statistic is a method used to quantify statistical heterogeneity in meta-analysis. It measures the percentage of total variation across studies that is due to actual differences (heterogeneity) rather than random chance. I² is calculated using the formula 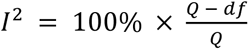, where Q is Cochran’s Q^73^ statistic and df is the degrees of freedom. Values range from 0% (no observed heterogeneity) to 100% (all variation is due to heterogeneity). Associations were interpreted as phylogenetically independent when they met three criteria: (i) meta-analytic significance (Stouffer’s p < 0.05), (ii) consistent effect direction across all phyla, and (iii) low heterogeneity (I² < 50%).

### Data Extraction Pipeline: Technical Implementation

Environmental data extraction was performed using the GEE JavaScript API via a multi-step pipeline. In coordinate preparation, sampling locations (decimal degree latitude/longitude) from 126 genome metadata records were formatted as an ee.FeatureCollection with geometry type ee.Geometry.Point. Coordinates underwent validation including range checks (-90° ≤ lat ≤ 90°, -180° ≤ lon ≤ 180°), ocean mask verification using the MODIS Land-Water Mask to identify coordinates erroneously falling on land pixels (n=3 adjusted to nearest ocean pixel within 5km), and duplicate detection flagging genomes sampled from identical locations. Image Collection Filtering and Temporal Aggregation: For each environmental variable, we loaded the corresponding GEE ImageCollection, applied temporal filters (2020-01-01 to 2023-12-31), selected the target band (e.g., ‘sst’ for temperature), and computed pixel-wise temporal statistics using ee.Reducer functions: var sstCollection = ee.ImageCollection(’NASA/OCEANDATA/MODIS-Aqua/L3SMI’) .filterDate(’2020-01-01’, ‘2023-12-31’) .select(’sst’); var sstMean = sstCollection.mean(); var sstMin = sstCollection.min(); var sstMax = sstCollection.max(); var sstStd = sstCollection.reduce(ee.Reducer.stdDev()). This produces four composite images where each pixel represents the temporal mean, minimum, maximum, or standard deviation across all valid observations in the 4-year window. Quality masking (excluding cloud-contaminated or sensor-flagged pixels) occurs automatically via built-in MODIS quality flags before aggregation.

For spatial sampling at point locations, temporal composite images were sampled at each genome’s coordinate using ee.Image.reduceRegion() with ee.Reducer.first() to extract the pixel value at the point location. We specified scale=1000 (1km) to approximate AEF resolution despite MODIS native 4km resolution; this uses nearest-neighbor resampling. Sensitivity analysis comparing 1km vs. 4km extraction had negligible differences (mean SST difference 0.03°C, r=0.998), confirming resampling does not introduce systematic bias. For nearshore samples (<5km from coast), we applied a 3 × 3 pixel median filter to reduce land contamination effects where ocean pixels mix with land in the sensor’s point spread function. The sampling operation was mapped across all features in the coordinate FeatureCollection using .map(), generating a new FeatureCollection where each feature contains original metadata plus extracted environmental values. Results were exported to Google Drive as CSV files via Export.table.toDrive().

### Quality Control for GEE Analyses

Satellite data quality varies spatially and temporally due to cloud cover, sun glint, sensor calibration drift, and atmospheric interference, necessitating systematic quality controls. We leveraged intrinsic quality flags embedded in MODIS products that indicate pixel-level retrieval confidence. For SST retrievals, we excluded pixels with quality_level > 2, while standard ocean color quality flags were applied to reject cloud-contaminated, ice-affected, or land-contaminated pixels. Temporal coverage requirements ensured environmental composites represented adequate observational periods rather than sparse, potentially unrepresentative subsets. Locations with < 50% valid observations after quality filtering were excluded; for SST daily products spanning 4 years (1,461 days), this required ≥730 valid days.

Five genomes (4%) failed this criterion, primarily from high-latitude regions (>60°N) with persistent cloud cover or seasonal sea ice (Table S1). Extracted values underwent physical plausibility checks with range validation applied for each variable: SST (-2°C to 35°C, spanning seawater freezing point to maximum tropical SST), chlorophyll-a (0.01 to 100 mg/m³, from oligotrophic gyres to extreme coastal blooms), and bathymetry (-11,000m to 0m). One spurious SST value (47.3°C) was detected and corrected by expanding the sampling region to a 3 × 3 pixel median filter to reduce potential land contamination artifacts.

### Derived Environmental Variables

Beyond directly extracted satellite variables, we computed derived metrics to capture key environmental gradients relevant to macroalgal physiology and adaptation. Thermal variability was quantified as the annual temperature range (sst_range = sst_max - sst_min), representing the full thermal amplitude experienced over the 4-year observation period and serving as a proxy for thermal tolerance requirements. Additionally, we calculated the temperature seasonality coefficient of variation (CV = sst_stddev / sst_mean) to provide a normalized variability measure enabling comparison across different thermal regimes. Chlorophyll-a concentrations were also categorized into standard oceanographic productivity classes—oligotrophic (<0.1 mg/m³), mesotrophic (0.1-1.0 mg/m³), and eutrophic (>1.0 mg/m³)—to facilitate categorical analyses of nutrient availability and primary productivity. The final GEE-derived environmental dataset comprised 126 genomes characterized by 13 environmental variables: sst_mean, sst_min, sst_max, sst_range, sst_stddev, chlor_a_mean, chlor_a_stddev, poc_mean, par_mean, bathymetry, distance_to_coast, latitude, and longitude.

### Vision Transformer [ViT] Application

AEF is a deep learning system developed by Google Research that encodes geographic locations into dense vector representations (embeddings) capturing multidimensional environmental context.^74^ The model architecture is based on ViT,^45^ a neural network design that processes images through self-attention mechanisms rather than traditional convolutional operations. AEF Foundations (AEF) is a geospatial foundation model that learns low-dimensional environmental embeddings directly from multi-spectral satellite imagery, rather than relying only on hand-crafted variables such as temperature or vegetation indices. AEF was trained at global scale on petabyte-scale imagery using multi-task learning objectives that include species-occurrence prediction from GBIF data, land-cover classification, climate-zone prediction, and vegetation-productivity proxies. These jointly trained tasks encourage the model to capture general-purpose environmental structure that can be reused across ecological applications We used the ViT architecture as it was originally configured. Input satellite images were divided into fixed-size patches (e.g., 16×16 pixels), each patch was linearly projected into a high-dimensional embedding space, positional encodings were added to preserve spatial relationships, the sequence of patch embeddings passed through multiple transformer encoder layers consisting of multi-head self-attention and feedforward networks, and the final hidden layer produced a fixed-length vector (64 dimensions for AEF) representing the environmental context of the location.

AEF embeddings are derived from the Sentinel-2 satellite constellation, operated by the European Space Agency as part of the Copernicus program.^75^ Sentinel-2 provides multispectral imagery at 10m spatial resolution in visible and near-infrared bands (blue, green, red, NIR), with additional bands at 20m (red-edge, short-wave infrared) and 60m (coastal aerosol, water vapor) resolution. The constellation consists of two satellites (Sentinel-2A launched 2015, Sentinel-2B launched 2017) in sun-synchronous orbits providing global coverage with 5-day revisit time at the equator. For coastal macroalgal habitats, 10m resolution captures fine-scale heterogeneity including: individual reef structures and substrate patterns (rocky vs. sandy), kelp forest canopy extent and patchiness, coastal development gradients, small-scale productivity hotspots (river plumes, upwelling zones), and wave exposure patterns (sheltered vs. exposed coastlines).

Spatial autocorrelation was assessed using Moran’s I ^76^ with inverse-distance weighting. To control for geographic confounding, we performed spatial regression including thin-plate spline basis functions of latitude and longitude (5 knots per dimension, including lat × lon interaction) as covariates, testing whether PFAM-environment associations remained significant after accounting for spatial location.

Some Pfams (PF12094: I=0.145, p=0.006; PF03537: I=0.128, p=0.018) do show spatial structure, which the phylogenetically-aware CV inherently controls for since phyla have biogeographic distributions. We note that the AEF training process integrates multi-year Sentinel-2 composites (2017-2024 for our analysis) to reduce noise from cloud cover, seasonal variation, and transient events, producing stable environmental representations reflecting long-term average conditions. The ViT architecture processes the full multispectral signature (all spectral bands simultaneously) rather than individual bands, enabling detection of environmental patterns visible only through spectral combinations—for example, distinguishing substrate types (rock vs. sand) through combined analysis of visible and short-wave infrared reflectance patterns invisible in single-band imagery.

### Interpreting the 64-Dimensional Embedding Space of AEF

The model’s final output is a 64-dimensional embedding vector for any geographic coordinate: embedding = f_AEF(lat, lon) ∈ ℝ⁶⁴ where each dimension (A00, A01, …, A63) is a continuous value typically ranging from -5 to +5 (mean ≈ 0 due to standardization during training). Unlike human-engineered features (temperature, chlorophyll), these dimensions are latent variables learned by the neural network and lack explicit physical interpretations. The dimensionality was selected via hyperparameter tuning during model development to balance expressiveness—sufficient capacity to encode complex environmental gradients globally, computational efficiency—low enough dimensionality for downstream machine learning tasks without excessive compute/memory requirements, and generalization—regularization to prevent overfitting to training data. Alternative dimensionalities (32, 128, 256) were tested, with 64 providing optimal performance on biodiversity prediction benchmarks.^74^ This is comparable to other representation learning systems (BERT-base: 768 dimensions, GPT-2: 1024 dimensions) but scaled for geographic data.

Associations between Pfam abundances and AEF geospatial embeddings were quantified using Spearman correlations across all 64 embedding dimensions and 10,300 Pfams (659,200 tests). Whether these clusters represent functionally coordinated modules or statistical groupings of individually associated varies by cluster. Statistical significance was assessed using Benjamini–Hochberg FDR correction (q < 0.05; see Tables S12-S16). Top associations were subsequently re-evaluated using the phylum-stratified framework described above to confirm lineage-independent signal when relevant. AEF satellite embeddings were extracted for 126 macroalgal genomes using the Google Earth Engine (GEE) Python API, accessing the precomputed annual ImageCollection (GOOGLE/SATELLITE_EMBEDDING/V1/ANNUAL). Sampling coordinates were first validated and converted to ee.Geometry.Point objects, matching the procedure used for environmental variable extraction to ensure methodologically consistent geolocation.

The AEF ImageCollection was mosaicked into a single 64-band image (bands A00–A63), representing the 2017–2024 annual composite. Embeddings were then retrieved in batch by applying reduceRegions with a mean reducer at 10-m resolution to a FeatureCollection containing all sampling locations. The PFAMs showing significant correlation with A06 (Table S10) were analyzed for domain-centric Gene Ontology (dcGO).^49^

To assess whether AEF vision transformer embeddings capture environmental gradients, we performed Spearman rank correlations between all 64 dimensions (A00 through A63) and two sets of environmental data: (1) collection-associated metadata (temperature, latitude, and longitude) across 126 macroalgal genomes, and (2) high-resolution GEE oceanographic features. Significance was assessed using the Benjamini-Hochberg method for manual metadata pairs and Bonferroni correction for GEE variables (p-values less than 6.0 × 10^5^).

### Correlation of Measured Oceanographic Variables with Learned AEF Embeddings

The 64 AEF dimensions capture comprehensive environmental gradients validated against independent Google Earth Engine (GEE) satellite products: 100% (64/64) dimensions correlated significantly (p < 0.05) with at least one GEE variable (sea surface temperature, depth, chlorophyll concentration, particulate organic carbon, distance from coast, or water clarity), with 67.2% (43/64) surviving Bonferroni correction. Sea surface temperature correlated with 58/64 dimensions (strongest: A52, r = -0.703), confirming that learned representations encode real oceanographic gradients rather than spurious satellite image artifacts.

AEF is a self-supervised vision transformer trained on petabyte-scale satellite imagery using objectives entirely independent of oceanographic measurements—specifically, species occurrence prediction, land cover classification, and climate zone identification. The model receives no sea surface temperature, chlorophyll, or bathymetry data during training. That the resulting 64-dimensional embeddings nevertheless correlate strongly with independently measured GEE variables (e.g., A52 with SST, r = −0.703) confirms that the learned representations encode genuine environmental structure rather than image artifacts, demonstrating that unsupervised representation learning recovers physically meaningful gradients without explicit supervision.

The two frameworks offer complementary strengths: GEE provides directly interpretable oceanographic parameters at 4-kilometer resolution (MODIS Aqua, Landsat)^77^, while AEF operates at 10-meter resolution (Sentinel-2) and integrates nonlinear, multivariate spectral patterns that may escape single-variable parameterization (e.g., potentially representing (reef structure, coastal geomorphology, small-scale habitat mosaics). Additionally, the 12 AEF dimensions uncorrelated with standard GEE variables (A11, A19, A20, A31, A34, A37, A38, A40, A55, A57, A58, A59) capture environmental features—such as seasonal thermal amplitude, coastal geomorphology, or productivity regimes—not represented in conventional oceanographic products. When Pfam associations converge across both frameworks, as observed for PF00092 (vWF-A), oxidative stress-related peroxidases, and metabolic enzymes, this concordance reflects shared underlying environmental signal rather than independent discovery, but it increases confidence that identified associations are robust to analytical approach and reduces the likelihood of methodological artifacts. Divergent associations, by contrast, identify candidates for novel environmental axes shaping macroalgal genomic composition.

The 100% agreement rate with GEE variables suggests that ViT can serve as “universal environmental sensors” capturing ocean conditions without requiring explicit physical measurements. This enables genomic prediction for unsampled locations based solely on satellite imagery—identifying likely genomic compositions of species inhabiting a given reef, predicting which genotypes are suited for translocation to degrading habitats, or forecasting range shifts under climate change by comparing genomic profiles to projected future imagery. However, interpretability remains a limitation: while dimension A52 correlates with temperature (r = -0.703), it is a latent variable without defined physical units, complicating mechanistic interpretation. Hybrid approaches combining learned embeddings with physically-interpreted variables may offer the best balance of predictive power and biological interpretability.

### Phylum-Environment Biogeography

Macroalgal genomes had phylum-level biogeographic structure as expected from their collection distributions. Ochrophyta (brown algae, n=43) predominantly occupy cold-water environments with a mean temperature of 12.2°C, representing 27.9% of polar samples but only 4.7% of tropical samples, and exhibiting the highest mean absolute latitude (45.5°) of the three phyla. In contrast, Rhodophyta are biased toward warmer waters (mean 17.2°C), with 27.1% occurring in tropical zones and only 12.9% in polar regions, while Chlorophyta (green algae, n=13) had moderate (mean 18.3°C). This biogeographic structuring creates a fundamental interpretive challenge: when Ochrophyta-specific Pfams correlate with cold-water AEF dimensions, does this reflect functional cold-adaptation of those protein domains, or simply the fact that Ochrophyta/Phaeophyta as a lineage happen to occupy cold environments? The within-phylum residuals correction addresses this by testing whether Pfams show environmental associations independent of phylum-level biogeography.

### Computation of Pfam–Environment Correlations

For each of 10,300 Pfam protein families, domain copy number was quantified across 126 macroalgal genomes, generating an abundance vector describing how many instances of that Pfam occur per genome. These vectors capture lineage-specific expansions and contractions of functional domains potentially linked to environmental adaptation. Each genome was then paired with its corresponding 64-dimensional AEF embedding, derived from satellite imagery at the collection coordinates using a self-supervised ViT. These latent dimensions (A00–A63) represent continuous environmental features learned directly from geospatial context, with values typically ranging from −0.5 to +0.5.

Pairwise Spearman rank correlations were computed between each Pfam’s abundance vector and each AEF dimension, producing a genome–environment association matrix. Spearman’s ρ was chosen over Pearson’s r because Pfam copy numbers are non-normally distributed, include zeros, and may relate to environmental gradients in a monotonic but nonlinear fashion. Each test yielded a correlation coefficient (ρ) and a p-value, with statistical significance evaluated using the Benjamini–Hochberg false discovery rate (FDR) procedure (pₐdⱼ < 0.05). The resulting matrix of Spearman coefficients encodes the direction and magnitude of association between protein family abundance and environmental embeddings.

Analysis of 2,022 significant Pfam-dimension associations (FDR < 0.05) revealed that most dimensions exhibit unidirectional effects: Pfam abundances consistently increase OR decrease along each environmental gradient, but not both, indicating that dimensions represent distinct, non-overlapping ecological axes rather than confounded variables. Network analysis revealed a hub-and-spoke architecture: dimension A25 connects to 49 Pfam domains (all negative correlations), while A18 connects to 48 domains (all positive), suggesting these represent opposite ends of a major environmental axis—potentially the cold coastal/upwelling gradient (A25) versus warm tropical/oligotrophic gradient (A18). This hub structure indicates that a relatively small number of environmental axes (the top 12 dimensions account for 78% of all associations) drive most of the genomic differentiation we observe, simplifying the search space for identifying climate-adapted genotypes.

### Bi-clustering of functional domains and AEF environmental embeddings

We quantified the relative abundance of 10,300 unique Pfam domains across all assembled macroalgal genomes and computed Pearson correlations with 64 AEF environmental embeddings per species. The embeddings were derived from satellite imagery using a self-supervised ViT trained to predict geographic coordinates from multispectral environmental context, independent of any biological labels. To enhance interpretability, each Pfam’s correlation profile was z-score normalized across all 64 dimensions, centering on its mean and scaling by its standard deviation, showing relative enrichment or depletion patterns rather than absolute magnitude. This z-scored correlation matrix was then subjected to bi-clustering to reveal coherent blocks of function–environment associations. AEF analysis identified significant associations between embedding dimensions and Pfam domain abundances (FDR < 0.05, Spearman |r| = 0.389 mean). Correlations were false discovery rate–corrected (Benjamini–Hochberg ^78^ FDR^78^ < 0.05) and visualized as a matrix of Pfams × AEF dimensions.

To identify coherent genomic–environmental associations, we applied iterative two-way hierarchical clustering (row-column bi-clustering) on the z-score–normalized correlation matrix, with normalization performed per Pfam (row-wise) to emphasize relative responses. This approach—analogous to row-centering in transcriptomics—enhanced detection of coordinated environmental responses among Pfams with otherwise low raw correlation magnitudes. Cluster structure was visualized in heatmaps using diverging color scales (blue for negative, tan/olive for positive). Latitude correlations for each AEF dimension were computed separately to assist in ecological interpretation of resulting clusters.

We performed phylo-aware and phylo-unaware clustering. The phylogenetic correction addresses a fundamental statistical problem in comparative genomics: phylogenetic non-independence. When we correlate Pfam abundances with environmental variables across 126 macroalgal genomes, standard statistical methods assume each genome is an independent observation. However, this assumption is violated because closely related species share genomic features due to common ancestry rather than independent environmental adaptation. In our dataset, the 126 samples are distributed across three major phyla—70 Rhodophyta (red algae), 43 Ochrophyta (brown algae), and 13 Chlorophyta (green algae)—and each phylum occupies distinct biogeographic regions with characteristic environmental conditions. This creates a confounding scenario: if Rhodophyta-specific Pfams are found predominantly in warm tropical waters, those Pfams will show strong correlations with “tropical” AEF embedding dimensions simply because of where red algae happen to live, not necessarily because those Pfams provide functional adaptations to tropical conditions.

To disentangle phylogenetic legacy from environmental adaptation, we applied a within-phylum residualization approach. For each Pfam domain, mean copy number was first calculated independently within each phylum. This phylum-specific mean was then subtracted from the Pfam count of each genome, yielding residual values that quantify deviations from the phylum average. This procedure removes between-phylum variance while preserving within-phylum variation.

Subsequent analyses correlated these residuals, rather than raw Pfam abundances, with AEF embedding dimensions. This framework addresses a distinct biological question: whether, within a given phylogenetic lineage, genomes exhibiting higher-than-average copy numbers of a given Pfam are associated with particular environmental conditions. Correlations that persist under this correction therefore reflect genuine within-lineage environmental associations, consistent with recent adaptive innovation, clade-specific functional evolution in response to local environments.

Several Pfams emerged with significant covariance across multiple analyses. PF01638 showed negative correlations with temperature in GEE phylo-ANCOVA (r=-0.446, p=9.48e-08), GEE Spearman (r=-0.489, p_fdr=2.42e-03), and per-phylum meta-analysis (pooled r=-0.483, p_fdr=1.03e-04, convergent across 3 phyla), plus associations with AEF dimensions A52 (r=0.450) and A53 (r=-0.472). PF12094 had the strongest temperature signal in GEE phylo-ANCOVA (r=-0.474, p=1.11e-08) and per-phylum meta-analysis (pooled r=-0.484, p_fdr=1.03e-04, convergent). PF00223 was not significant for environmental variables but showed associations with 9 AEF dimensions, most strongly A17 (r=-0.476, p_fdr=5.14e-03) and A36 (r=0.439, p_fdr=1.34e-02). PF14450 correlated negatively with temperature in GEE phylo-ANCOVA (r=-0.383, p=6.23e-06), per-phylum meta-analysis (pooled r=-0.425, p_fdr=2.00e-03, convergent), and AEF dimension A00 (r=-0.448, p_fdr=9.40e-03).

### Per-Phylum Bi-clustering Analysis

Within-phylum Pfam-AEF correlation matrices were computed separately for Rhodophyta (n=70), Ochrophyta (n=43), and Chlorophyta (n=13) using Spearman correlation. Pfams present in fewer than 3 samples per phylum were excluded. Significance was assessed using permutation testing (10,000 permutations) with Benjamini-Hochberg FDR correction. For Rhodophyta, bi-clustering was performed using two methods: (1) Spectral bi-clustering via scikit-learn’s SpectralBi-clustering (n_clusters=(20, 8), method=’log’, random_state=42), and (2) K-means bi-clustering combining hierarchical clustering of Pfams with K-means clustering of dimensions (n_clusters=20 for Pfams, n_clusters=8 for dimensions). Bi-cluster statistics (mean correlation, standard deviation, number of significant associations) were computed for each row-column cluster intersection. Method agreement was assessed using Adjusted Rand Index.

For bi-clustering analysis, we applied a stricter PFAM inclusion criterion (present in ≥10 samples) compared to the cross-phylum comparison (present in ≥15% of samples), reducing the Rhodophyta dataset from 6,953 to 4,747 PFAMs. This filter ensures robust correlation estimates by excluding sparsely represented domains while reducing the multiple testing burden, as FDR correction severity scales with the number of tests performed. Functional annotations for Pfams in top bi-clusters were retrieved from the InterPro API (https://www.ebi.ac.uk/interpro/api/entry/pfam/) and manually categorized into functional groups.

### Domain-Centric Gene Ontology (dcGO) Enrichment Analysis

To identify functional enrichment among Pfam domains associated with specific environmental categories, we performed domain-centric dcGO enrichment analysis using the dcGO database. Unlike conventional gene-level enrichment tools (e.g., DAVID), dcGO maps ontology annotations directly to protein domains, enabling functional inference for domain lists derived from comparative genomics without requiring full-length gene identifiers. For each environmental comparison, we compiled Pfam domain lists meeting two criteria: (1) unique presence in the target environmental category (e.g., saltwater versus freshwater species), and (2) significant correlation with AEF embedding dimensions (FDR < 0.05). These domain lists were submitted to the dcGO enrichment facility (http://www.protdomainonto.pro/dcGO) with the following parameters: Domain classification: Pfam domains were used as the input domain type, leveraging dcGO’s pre-computed Pfam-to-GO mappings derived from UniProt annotations propagated through the GO-directed acyclic graph (DAG) structure following the true-path rule.

Enrichment significance was assessed using the hypergeometric distribution, which models the probability of observing k or more domains annotated to a given GO term in a sample of n domains drawn without replacement from a background of N annotatable domains containing K domains with that annotation: P(X ≥ k) = Σ [C(K,i) × C(N-K, n-i)] / C(N,n) for i = k to min(n,K) where C denotes the binomial coefficient. The hypergeometric test was selected over Fisher’s exact test for increased sensitivity in detecting enriched terms (https://www.rdocumentation.org/packages/dcGOR/versions/1.0.6/topics/dcEnrichment). All Pfam domains with at least one GO annotation in the dcGO database served as the background population.

For multiple testing correction, P-values were adjusted using the Benjamini-Hochberg false discovery rate (FDR) procedure, with significance declared at FDR < 0.05. Z-scores were computed based on the deviation of observed overlap from expected overlap under the null hypothesis. Ontology categories: Enrichment was performed separately for three GO categories—Biological Process (BP), Molecular Function (MF), and Cellular Component (CC)—to distinguish functional roles, biochemical activities, and subcellular localization patterns among environment-associated domains.

## ACKNOWLEDGEMENTS

We would like to thank the NYUAD sequencing core of the Core Technologies Platform (CTP), the NYUAD Bioinformatics core, and the NYU (Greene) and NYUAD (Jubail) HPC platforms. This research was supported by NYUAD Faculty Research Funds (AD060) and Tamkeen under the NYU Abu Dhabi Research Institute Award to the NYUAD Center for Genomics and Systems Biology (73 71210 CGSB9).

## AUTHOR CONTRIBUTIONS

K.S.-A. and A.M. conceptualized the study. A.M., N.A.-M. and J.A.B. collected the field macroalgal samples. L.N. took and edited the herbarium photos. C.R.-M., M.S. and D.H.G. extracted DNA. K.S.-A., A.M., D.C.E.-A., A.K.J, and D.R.N. designed computational data analysis. A.M., K.S.-A., D.C.E.-A., A.K.J., D.R.N., and N.D. performed computational data analysis. A.M., D.R.N., S.T.E., and M.A.-H. did statistical data analysis. A.M, D.C.E.-A. and D.R.N. prepared the figures. A.M., K.S.-A., D.R.N., and D.C.E.-A. wrote the manuscript with input from all authors. K.S.-A. supervised the study.

## DECLARATION OF GENERATIVE AI AND AI-ASSISTED TECHNOLOGIES IN THE WRITING PROCESS

The authors used several AI models in the writing process. After using this tool/service, the authors reviewed and edited the content as needed and take full responsibility for the content of the publication.

## RESOURCE AVAILABILITY

### Lead contact

Further information and requests for resources and reagents should be directed to and will be fulfilled by the lead contact, Kourosh Salehi-Ashtiani.

### Data availability statement

The raw reads of the assemblies are in NCBI BioProject: PRJNA929663. All the data and code used in data processing are available online accompanying this manuscript and in Zenodo repositories (doi: 10.5281/zenodo.17775003 (https://zenodo.org/records/18051219)) as described in the supplementary materials. Macroalgae assemblies (original and decontaminated) can be found in Zenodo (doi:10.5281/zenodo.7758508).

### Code availability statement

The programs and scripts used in this work are available in Data S2 which is available in the supplementary materials accompanying this manuscript and also at Zenodo (doi: 10.5281/zenodo.17775003 (https://zenodo.org/records/18051219)).

## SUPPLEMENTAL INFORMATION

### SUPPLEMENTAL TABLES

**Table S1. Genome Assembly and PFAM Annotation Data**

Table S1A. New Species Metadata. This table contains metadata for 9 newly sequenced macroalgal genomes from the Arabian Gulf region. Data include collection information (date, geographic coordinates, depth, temperature), assembly metrics (N50 values at different k-mer sizes, output in GB), and transcriptome quality assessment (BLEACHed tORFeomes counts and BUSCO completeness scores across two rounds of analysis). Species represented include Padina boergesenii, Polycladia myrica, Avrainvillea amadelpha, and others from Phaeophyta, Chlorophyta, Rhodophyta.

Table S1B. Assembly Benchmarking. Comparison of de novo assembly performance across three assemblers (CLC Genomics Workbench, ABySS, and SPAdes) using multiple k-mer sizes (40, 50, 60). N50 values are reported for each combination to identify optimal assembly parameters.

Table S1C. Phylogeny. Taxonomic classification for the macroalgal genomes including species, family, order, class, and phylum assignments. Morphological classifications and rbcL gene sequences are provided for phylogenetic reconstruction. The dataset spans 3 phyla (Phaeophyta, Chlorophyta, Rhodophyta) and 8 classes.

Table S1D. Reference Genomes. Published reference genomes included in this study with associated metadata. Columns include genome identifiers, literature citations, species names, salinity regime, climatic zone, geographic location, temperature, coordinates, and habitat type.

Table S1E. PFAM Counts with Metadata. PFAM protein domain counts for 131 macroalgal genomes across 10,589 domain families, which was filtered to 10,300 PFAMs to remove all-zero and invariant domains and 126 genomes to include only those samples with extractable GEE data. Each row represents a genome with associated metadata columns (species, phylum, climatic zone, temperature, habitat, coordinates) followed by count values for each PFAM domain identified via hmmsearch against the Pfam-A database.

**Table S2. Google Earth Engine Environmental Correlation Analysis**

Table S2A. Environmental Variables per Genome. Environmental parameters extracted from Google Earth Engine. Variables include sea surface temperature statistics (mean, min, max, range), chlorophyll-a concentration (mean, standard deviation), particulate organic carbon, bathymetry, and distance to coast. Climate zones represented: Polar, Subpolar, Subtropical, Temperate, Tropical.

Habitats: subtidal, intertidal, coralligenous, epiphyte, endophyte, mangrove, natural_pool_, salt_marsh, seagrass, spring, aquarium, oyster.

Table S2B. GEE Phylogenetically-Corrected ANCOVA. Results of phylogenetically-corrected analysis of covariance (ANCOVA) testing associations between 338 PFAM domains and environmental variables. Columns include PFAM accession, environmental variable, overall correlation coefficient (r), p-value, Stouffer combined p-value, heterogeneity metrics (I², Q statistic), number of phyla tested, and per-phylum correlation coefficients for Rhodophyta, Ochrophyta, and Chlorophyta.

Table S2C. All GEE Correlations. Complete Spearman correlation results for 78,143 PFAM-environment pairs across 6,011 unique domains and 13 environmental variables. Columns include correlation coefficient, raw p-value, sample size, prevalence (proportion of genomes with non-zero counts), direction of effect, Bonferroni-corrected p-value, FDR-adjusted p-value, and significance flags. 15 associations were significant at FDR<0.05; 7 passed Bonferroni correction.

Table S2D. Per-Phylum Meta-Analysis. Meta-analysis of PFAM-environment correlations performed separately within each phylum and combined using Stouffer’s Z-score method (16,047 total tests). Columns include pooled correlation coefficient, Stouffer Z and p-values, heterogeneity statistics, directionality metrics, and per-phylum correlations with p-values. Interpretation categories indicate whether effects were CONVERGENT (same direction across phyla), DIVERGENT (opposite directions), or NOT_SIGNIFICANT. 2,431 associations showed convergent patterns.

**Table S3. AlphaEarth Analysis**

Table S3A. PFAM-Embedding Dimension Associations. Significant correlations between PFAM domain abundances and AlphaEarth embedding dimensions. The AlphaEarth framework projects genomes into a learned embedding space where dimensions capture environmental gradients.

Columns include PFAM accession, embedding dimension, Pearson correlation, raw p-value, sample size, and FDR-adjusted p-value. 313 associations were significant at FDR<0.05.

Table S3B. Phylogenetically-Corrected PFAM-Embedding Associations. Phylogenetically-corrected correlations between PFAM domains and AlphaEarth embedding dimensions (843 tests).

Phylogenetic correction accounts for shared evolutionary history that could confound environment-genome relationships. 843 associations remained significant at FDR<0.05 after phylogenetic correction.

Table S3C. Embedding Dimension-Environment Correlations. Correlations between AlphaEarth embedding dimensions and environmental variables. This table validates that embedding dimensions capture meaningful environmental gradients by testing their associations with measured variables including Latitude, Longitude, Climatic zone, Habitat, Phylum. Both Pearson and Spearman correlations are reported with FDR-adjusted p-values.

Table S3D. Embedding Dimension Rankings. Summary rankings of AlphaEarth embedding dimensions by the number of significantly associated PFAM domains. Columns include dimension identifier, count of significant PFAM associations (FDR<0.05), the strongest associated PFAM, its correlation coefficient, and the number of significant GEE environmental correlations for that dimension. The top-ranked dimension was A25 with 49 significant PFAM associations.

### SUPPLEMENTAL DATA

**Data S1. RbcL phylogenetic analysis files.** Archive containing phylogenetic reconstruction data for macroalgal classification. Contents: (1) rbcL_alignment.fa - MUSCLE-aligned amino acid sequences of ribulose-1,5-bisphosphate carboxylase/oxygenase large subunit (RbcL) markers extracted from 119 macroalgal genomes using tBLASTn searches with phylum-specific queries (Ectocarpus sp. AAR24444.1 for Ochrophyta, Gracilariopsis hommersandii AAL46528.1 for Rhodophyta, Ulva rigida QWL15241.1 for Chlorophyta); (2) rbcL_phylogenetic_tree.nwk - Maximum likelihood phylogenetic tree in Newick format, reconstructed using MEGA v11 with Jones-Taylor-Thornton (JTT) substitution model, gamma-distributed rates (4 categories), Nearest-Neighbor-Interchange heuristic search, and 1000 bootstrap replicates. Tree visualization performed in iTOL v6. File format: gzip-compressed tar archive (.tar.gz).

**Data S2. Tarball archive containing the scripts used to perform the analyses reported in this study.** All the data and code used in data processing are available online accompanying this manuscript and in Zenodo repositories (doi: 10.5281/zenodo.17775003 (https://zenodo.org/records/18051219)).

